# Common physiological processes control mercury reduction during photosynthesis and fermentation

**DOI:** 10.1101/2020.10.08.332528

**Authors:** Daniel S. Grégoire, Sarah E. Janssen, Noémie C. Lavoie, Michael T. Tate, Alexandre J. Poulain

**Author notes:** Corresponding author: Dr. Daniel S. Grégoire. Biology Department, University of Waterloo, 200 University Ave W, Waterloo, ON, N2L 3G1.

## Abstract

Mercury (Hg) is a global pollutant and potent neurotoxin that bioaccumulates in food webs as monomethylmercury (MeHg). The production of MeHg is driven by anaerobic and Hg redox cycling pathways such as Hg reduction, which control the availability of Hg to methylators. Anaerobes play an important role in Hg reduction in methylation hotspots, yet their contributions remain underappreciated due to how challenging these pathways are to study in the absence of dedicated genetic targets and low levels of Hg^0^ in anoxic environments. In this study we used Hg stable isotope fractionation to explore Hg reduction during anoxygenic photosynthesis and fermentation in the model anaerobe *Heliobacterium modesticaldum* Ice1. We show that cells preferentially reduce lighter Hg isotopes in both metabolisms leading to mass-dependent fractionation, but mass-independent fractionation commonly induced by UV-visible light is absent. We show that isotope fractionation is affected by the interplay between pathways controlling Hg recruitment, accessibility, and availability alongside metabolic redox reactions. The combined contributions of these processes lead to isotopic enrichment during anoxygenic photosynthesis that is in between the values reported for anaerobic respiratory microbial Hg reduction and abiotic photoreduction. Isotope enrichment during fermentation is closer to what has been observed in aerobic bacteria that reduce Hg through dedicated detoxification pathways. These results demonstrate that common controls exist at the atomic level for Hg reduction during photosynthesis and fermentation in *H. modesticaldum*. Our work suggests that similar controls likely underpin diverse microbe-mediated Hg transformations that affect Hg’s fate in oxic and anoxic habitats.

**IMPORTANCE:** Anaerobic and photosynthetic bacteria that reduce mercury affect mercury delivery to microbes in methylation sites that drive bioaccumulation in food webs. Anaerobic mercury reduction pathways remain underappreciated in the current view of the global mercury cycle because they are challenging to study, bearing no dedicated genetic targets to establish physiological mechanisms. In this study we used stable isotopes to show that common physiological processes control mercury reduction during photosynthesis and fermentation in the model anaerobe *Heliobacterium modesticaldum* Ice1. The sensitivity of isotope analyses highlighted the subtle contribution of mercury uptake towards the isotope signature associated with anaerobic mercury reduction. When considered alongside the isotope signatures associated with microbial pathways for which genetic determinants have been identified, our findings underscore the narrow range of isotope enrichment that is characteristic of microbial mercury transformations. This suggests that there exist common atomic-level controls for biological mercury transformations across a broad range of geochemical conditions.

## INTRODUCTION

Mercury (Hg) is a global pollutant and potent neurotoxin that bioaccumulates in aquatic and terrestrial food webs as monomethylmercury (MeHg) (1). Anaerobic microbes with the *hgcAB* gene cluster, which encodes for the metabolic machinery responsible for Hg methylation (2), contribute to MeHg production in habitats such as aquatic sediments, water columns, and flooded soils (3–6). Hg methylation is thought to occur intracellularly and as such, it is ultimately controlled by the bioavailability of Hg to methylators in anoxic habitats (7).

Hg redox cycling plays a key role in determining the inorganic Hg substrates available to methylators. Its role is two-fold: anaerobic Hg^0^ oxidation can supply dissolved Hg^II^ required to generate MeHg and evasion of Hg^0^ due to reduction can limit MeHg production by removing inorganic Hg substrates. Although it is well established that abiotic photochemical Hg reduction dominates Hg redox cycling in oxic surface systems where light is present (8–16), Hg reduction pathways in anoxic zones where Hg methylation occurs have mostly been characterized under laboratory conditions and remain challenging to assess in the field.

Anoxic Hg reduction can occur via abiotic redox reactions with dissolved organic carbon (17) and magnetite (18, 19), but also through biotic pathways mediated by chemotrophic and phototrophic microorganisms (20–24). Unlike many aerobes that reduce Hg using dedicated enzymatic machinery encoded by the *mer* operon (25, 26), Hg reduction in anaerobes occurs through cometabolic pathways tied to anaerobic respiration, fermentation, and anoxygenic photosynthesis (20–24). In contrast to *mer*-mediated Hg reduction and demethylation, dedicated genetic determinants have yet to be identified for anaerobic Hg reduction, making these pathways challenging to study from a mechanistic standpoint. Furthermore, while some studies have shown that Hg^0^ can significantly contribute to Hg speciation in anoxic habitats (27, 28), it is possible that low levels of Hg^0^ are maintained by an active cycle of reduction and oxidation (29). In the absence of genetic targets and measurable Hg^0^ accumulation, the contributions of anaerobic Hg^0^ production to global Hg cycling remain cryptic and difficult to assess.

The use of stable Hg isotope fractionation is a powerful tool for providing biogeochemical proxies for different abiotic and biotic Hg transformations (30, 31). All abiotic and biotic Hg transformation pathways studied to date demonstrate kinetic mass dependent fractionation (MDF) in which the product pool becomes isotopically lighter (enrichment of light isotopes) as the reaction proceeds (32). Some pathways, such as abiotic photochemical reduction and demethylation, also demonstrate mass independent fractionation (MIF), which is defined as the fractionation that occurs outside of the predicted MDF pattern (30). MIF is most commonly observed due to the magnetic isotope effect for odd-Hg isotopes ^199^Hg and ^201^Hg in the presence of sunlight (33) and the extent to which MIF occurs is controlled in part by water chemistry and the presence of photosynthetic organisms (39).

The combination of MDF and MIF tracers, and the rate of isotopic enrichment, referred to as fractionation factor, can be used to outline the underlying components of different Hg microbe-mediated Hg transformation pathways. Over the last decade, Hg isotope fractionation has been studied in microbes capable of *mer*-mediated reduction (34) and MeHg demethylation (35); anaerobic non-*mer*-mediated Hg reduction (36, 37); Hg methylation (37–40); and oxygenic phototrophic MeHg degradation and Hg reduction (41). These studies have yielded valuable insights and mechanistic details that explain how microbes interact with Hg at the atomic level making them a promising option for studying mechanisms for anaerobic Hg reduction.

Our objective in this study was to characterize Hg isotope fractionation patterns during anoxygenic phototrophic and fermentative Hg^II^ reduction, for which there exists no data. By providing the first isotopic characterization of these pathways we aim to better understand the underlying physiological pathways involved in anaerobic Hg reduction and their role in controlling Hg availability to methylators in anoxic habitats. In this work, we test whether Hg reduction by the model anaerobe *Heliobacterium modesticaldum* Ice1 during anoxygenic photosynthesis and fermentation leads to distinct isotope enrichment signatures, which would suggest the different metabolic Hg reduction pathways are controlled by separate processes. We also compare our results to previously published isotope signatures for biotic and abiotic Hg transformations to critically evaluate whether common controls exist for microbe-mediated Hg transformations. Finally, we discuss whether current analytical capacity for isotope fractionation can be used to track the contributions of such pathways in the environment.

## METHODS

### Cell growth conditions and bioreactor setup

All experiments with live cells employed the strain *Heliobacterium modesticaldum* Ice1 grown in PYE medium. Phototrophic cultures were grown at a constant visible light intensity of 80 μmol photon m^−2^ s^−1^ with a peak irradiance in the near-red to red spectrum (600 nm to 700 nm) at 50°C whereas chemotrophic cultures were grown in the dark at 50°C in line with our previous work (23). *H. modesticaldum* Ice1 was obtained from the DSMZ culture collection (catalogue number DSM-792). The bioreactor methodology employed in these experiments was similar to the one used in previous work (23) with the following modifications: the bioreactor was periodically supplied with sterile nitrogen gas that passed through an activated carbon filter and a 0.2 μm pore size air filter at a flow rate of 0.25 L min^−1^ and two gold traps were connected in series at the reactor outlet to capture Hg^0^ (**Fig S1**).

The bioreactor was kept in an incubator set to 50°C. Phototrophic experiments were performed using a 60 W incandescent light bulb with a visible light intensity being maintained at 20 μmol photon m^−2^s^−1^ and mirroring the irradiance of growth conditions. Chemotrophic experiments were performed in the dark. Background Hg^0^ was purged and removed from reactors prior to phototrophically or chemotrophically-grown *H. modesticaldum* cells being added as a 10 % (v/v) inoculum wherein an additional subsample was withdrawn to verify initial cell density via O.D. 600 nm. For sterile treatments, no cells were added to the bioreactor and the volume of growth medium was adjusted to ensure experiments were carried out at a working volume of 500 mL. In all experiments, NIST 3133, a standard with known Hg isotopic composition, was added to the bioreactor to approximately10 nM Hg final concentration. Additional details on bioreactor setup are provided in **Supporting Methods**.

### Hg^0^ measurements

The bioreactor was bubbled periodically rather than continuously in contrast to previous work because continuous bubbling led to fouling of gold traps and poor total Hg (THg) and isotope recoveries. Subsamples were collected at designated timepoints by connecting dual gold traps to the reactor, purging the reactor for 30 min to collect Hg^0^. In the final 5 mins the reactor liquid was sampled for Hg^II^ concentration, isotopic composition, and cell growth after which gas flow was stopped. All gold traps were capped to prevent Hg^0^ loss and aqueous samples were conserved with 1% (v/v) trace element grade HCl at 4°C in the dark. Reactor washes were performed following previously established protocols (23).

### Total Hg analyses and sample preparation for stable isotope measurements

All samples for aqueous Hg analyses were oxidized with 10 % (v/v) bromine chloride (BrCl) in line with previous work (40) and THg concentrations were determined by stannous chloride reduction coupled to cold vapor atomic fluorescence spectrometry (42). During THg analysis, duplicates and matrix standard spikes were analyzed every 10 samples; analyses showed less than 10 % relative percent difference between duplicates and spike recoveries of 90-110%. The detection limit was 0.2 pM.

Total Hg mass balances were calculated in line with previous work (23, 24) with corrections to account for the mass of Hg removed for each subsample. For Hg^0^ collected onto gold traps, a thermal desorption system was used to liberate Hg^0^ over a 40 min desorption cycle which was then captured in 40% HNO_3_:BrCl (3:1) oxidizing solution. An aliquot of the trapping solution was subsequently analyzed for THg concentration following the protocol established in previous work (43). Desorption efficiency during pre-concentration was tested by trapping a NIST 3133 standard with each batch of samples, and recoveries for standard trapping were 101.27 ± 4.26 (n = 15). THg recoveries from bioreactor experiments were 94.90 ± 9.50 % (n=7) (**Fig S2**).

### Hg stable isotope analyses

Using the previously determined THg concentration, an aliquot of the trapping solution was diluted with ultra-high purity water to an acid content < 10% H^+^ and a THg concentration of 3.75 to 5.00 nM (44). A Neptune Plus multicollector-inductively coupled plasma mass spectrometer (MC-ICP-MS) was used for Hg stable isotope ratio measurements. A concentration and matrix-matched NIST 3133 Hg was used for standard sample bracketing (45). To achieve detection at low Hg concentrations, Hg was reduced with stannous chloride in-line then introduced continuously into a custom designed gas liquid separator along with a thallium standard (NIST SRM 997, 195 nM in 3% HCl) for mass bias correction (44, 46). MC-ICP-MS instrument conditions and analytical expectations derived from previously published work and quality control metrics for Hg stable isotope analysis are provided in **Supporting Methods, Table S1** and **Table S2**.

### Isotope calculations

Delta calculations followed the conventions set forth by others (45). This convention calls for mass dependent fractionation (MDF) to be expressed in terms of δ^xxx^Hg where graphically δ^202^Hg is used to signify MDF. δ^xxx^Hg is calculated as:

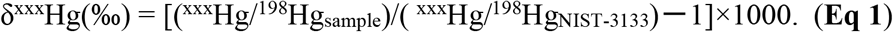

XXX is used to signify the isotope of interest. Hg also undergoes MIF of both even and odd isotopes. Here, odd-MIF is described by Δ^199^Hg and even-MIF by Δ^200^Hg. MIF is calculated as:

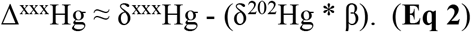

β denotes the mass dependent scaling constant, which is determined by the laws of mass dependence (45). Isotope enrichment effects based on the ratio of products to reactants (i.e. ε_p/r_, herein referred to as ε_AP_ for anoxygenic phototrophic reduction and ε_FM_ for fermentative reduction) were calculated using Rayleigh fractionation models to account for the Hg^0^ that was removed following methods from previous work on microbial Hg reduction (34).

This method involved fitting a linear regression (shown in **Fig S3, Fig S4**, and **Table S3**) based on the change in relative isotope ratios, ln (R/R_0_), as a function of ln(f_r_) where:

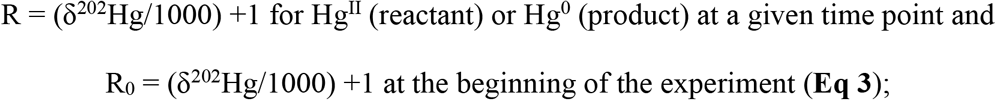

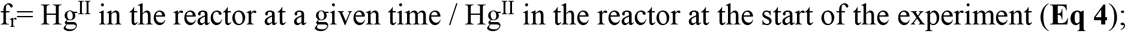

and the slope of the linear regression, which is equivalent to ε_AP_ or ε_FM_:

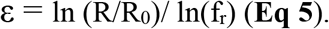

We chose to use the total Hg analyses from the bioreactor to establish f_r_ as this is a true representation of the instantaneous isotope fractionation rather than a time integrated sample following the methods outlined in previous work (34). Note that in our slope calculations we do not have data for ^202^Hg^0^ when f_r_=1 (the 0h sampling point) because no isotope fractionation was taking place prior to the start of Hg reduction (**Table S2** and **Table S3**). Additional calculations for isotope mass balances were carried out to test for deviation from the Rayleigh fraction model and can be found in **Table S4**. ε values for MDF were compiled from previous studies on pure microbial cultures and abiotic Hg transformations for comparison purposes. Currently, there is no standard guideline for how to calculate ε and studies vary widely in how they present isotope enrichment data and their associated variability. We have provided details on how we normalized ε values from different studies in **Table S5**. In this work we only compared enrichment factors from studies that used MC-ICP-MS techniques that were in line with our analyses, however we have included all of the enrichment factors for microbe-driven Hg transformations published to date in **Table S5**.

## RESULTS AND DISCUSSION

### Hg^0^ production during anoxygenic photosynthesis and fermentation

In our experiments Hg^0^ production relied on the presence of live cells for both chemotrophic and phototrophic growth conditions. Cumulative Hg^0^ production was an order of magnitude higher for live cell treatments compared to abiotic controls (>0.60 vs <0.05 nmol) (**Fig 1**). Chemotrophic cells produced double the amount of Hg^0^ compared to phototrophic cells (1.33 to 1.64 nmol vs 0.60 to 0.81 nmol, respectively) despite having been supplied with slightly less Hg^II^ (**Fig 1B, E**). In relative terms, phototrophic cells reduced 12 to 15% of the initial Hg^II^ supplied whereas chemotrophic cells reduced 31 to 40% (**Fig S2**).

**Figure 1:**
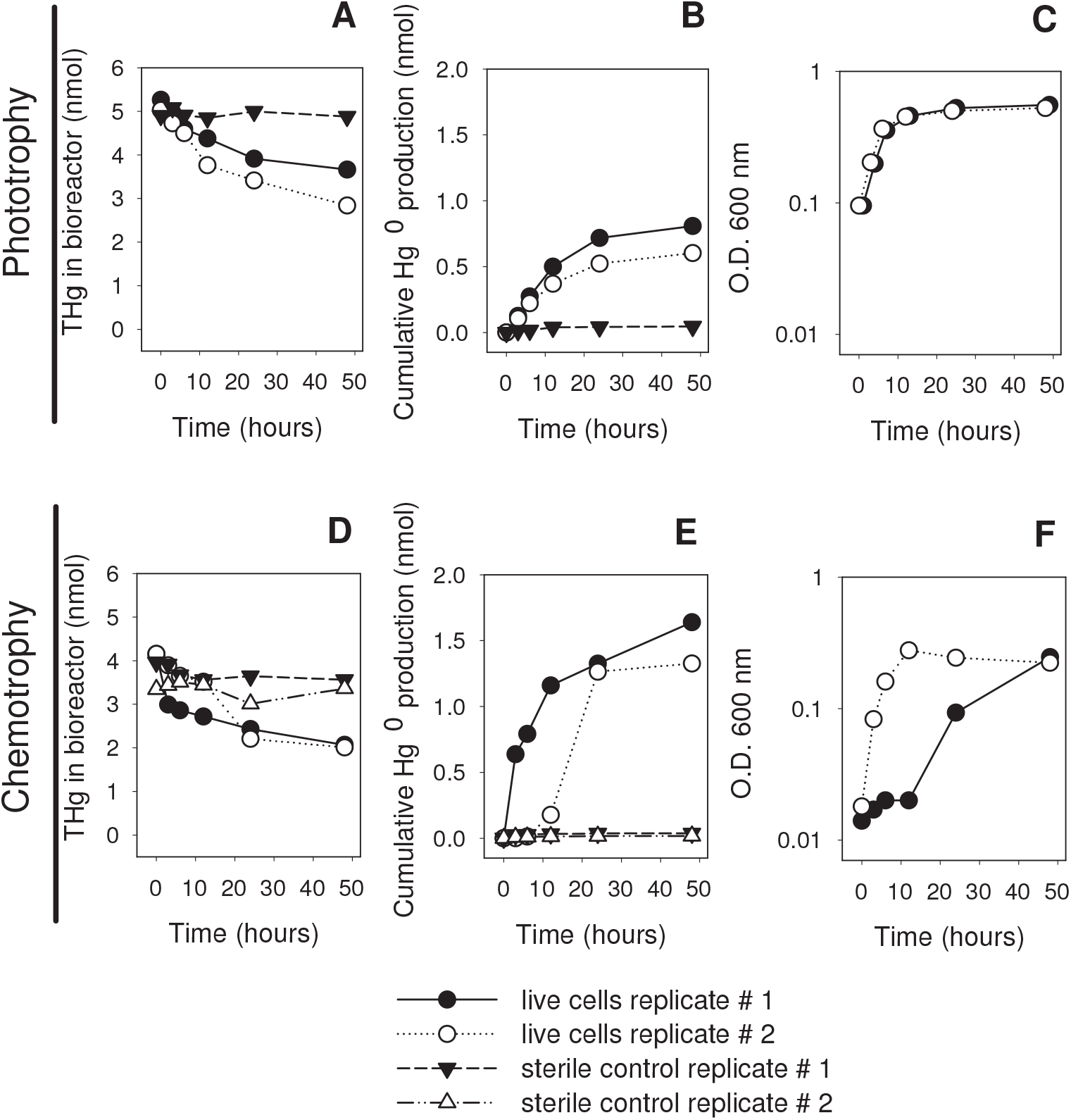
Total Hg and cumulative Hg^0^ production by *H. modesticaldum* grown phototrophically and chemotrophically. **A, D** THg for live cells and sterile controls without cells. **B, E** Cumulative Hg^0^ production for live cells and sterile controls without cells. **C, F** Microbial growth as measured by O.D. 600 nm for live cells and sterile controls without cells. 10 nM Hg was supplied in all experiments from the same stock (NIST 3133).

Relative Hg^0^ production in these experiments was lower, yet consistent with our previous work, which showed chemotrophically-grown cells reduced 15% more Hg^II^ than phototrophically-grown cells, reducing between 60 to 75% of the initial Hg^II^ supplied (23). The relative amount of Hg^0^ production did not increase with higher Hg^II^ exposure (10 nM) versus previous experiments (250 pM) (23). It is unlikely that the lower reduction rate observed in the experiments presented here was due to the toxicity of 10 nM Hg supplied, given that *H. modesticaldum* grows well under the same conditions at 50 times higher Hg concentrations (23). Instead, we suspect that the lack of constant bubbling of the reactor may be responsible for the observed results. It is possible that the accumulation of CO_2_ in the bioreactor following organic carbon oxidation may compete with Hg^II^ as an electron sink, decreasing the amount of Hg^0^ produced. Inorganic carbon is required for anaplerotic reactions that fulfill the biosynthetic needs of *H. modesticaldum* and CO_2_ could have competed with Hg^II^ for reducing power, as previously shown (47). These results could also be due to Hg^0^ oxidation allowed by the increased residence time of Hg^0^ in the reactor afforded by the intermittent sparging, as has been observed in other anaerobes (48, 49).

Despite carefully controlling the growth conditions in our experiments, chemotrophic cells from replicate #1 exhibited a lag phase of 12 hours (**Fig 1F**). Although this lag did not affect final cell density, the slow-growing culture from replicate #1 initiated Hg^0^ production earlier than the culture in replicate #2 (**Fig 1E, F**). These results may be attributable to cells in replicate #1 maintaining a higher Hg to cell ratio in the first 12 hours of the experiment, which suggests that Hg is being reduced intracellularly in line with our previous work on Heliobacteria (23, 50). Indeed, even at low initial densities, cells can maintain μM levels of intracellular reducing power, which could be used to reduce nM levels of Hg^II^ (24).

### Mass dependent fractionation during photosynthetic and fermentative growth

Hg reduction in phototrophically grown *H. modesticaldum* resulted in consistent positive MDF whereas fermentatively grown cultures showed positive MDF with variability in fractionation patterns across replicates (**Fig 2**). Abiotic controls showed minimal fractionation for both MDF and MIF, confirming that Hg reduction and subsequent isotopic fractionation were predominantly driven by cellular processes (**Fig 2**). Live cell experiments were devoid of MIF (**Table S1** and **Table S2**).

**Figure 2:**
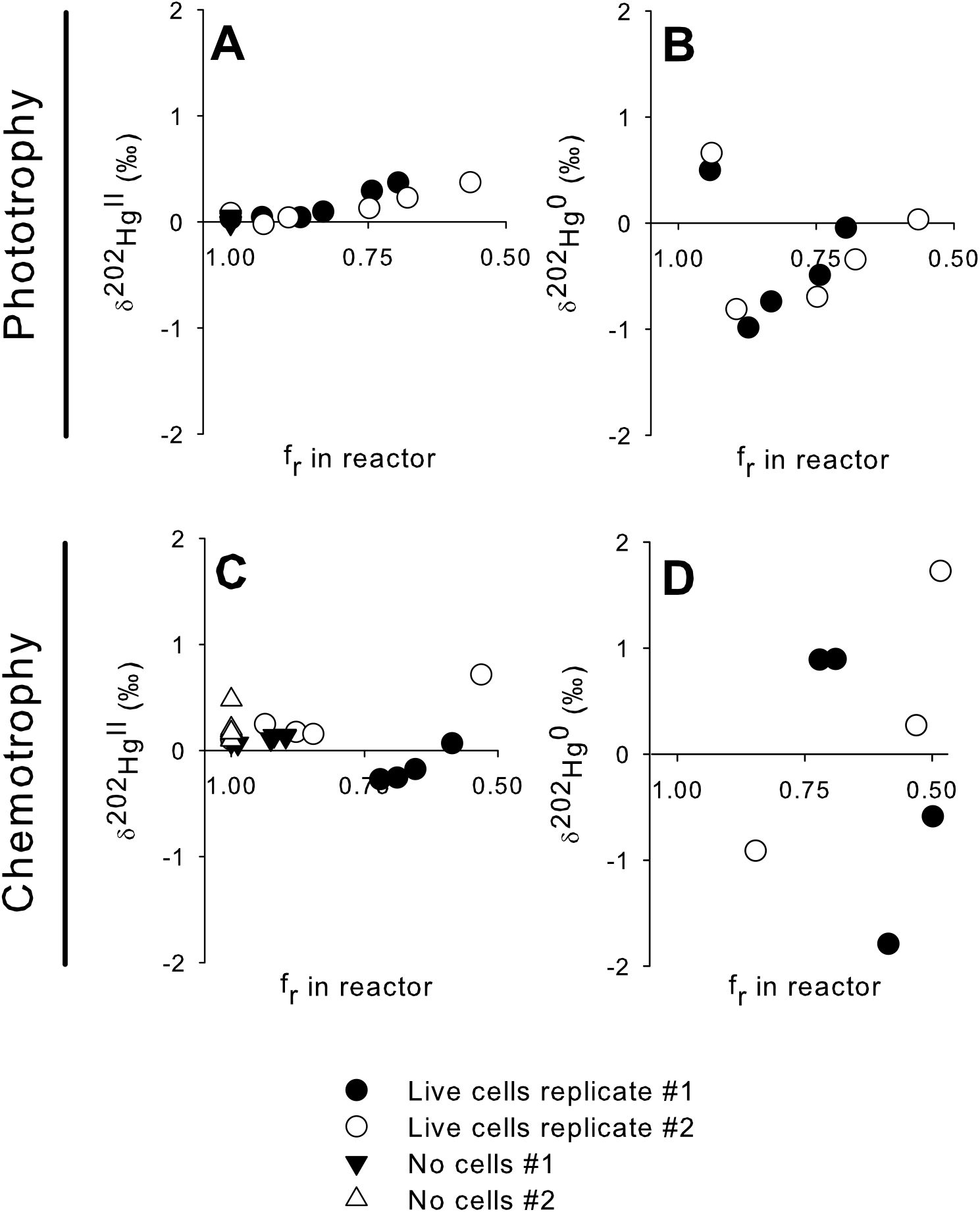
Mass dependent fractionation of ^202^Hg in Hg^II^ (A, C) and Hg^0^ (B, D) during phototrophic and chemotrophic growth of *H. modesticaldum* and no cell controls. δ^202^Hg values for Hg^II^ and Hg^0^ are plotted with respect to the fraction of remaining total inorganic Hg^II^ in the bioreactor. Insufficient Hg^0^ was recovered from the no cell controls and thus no results are displayed. 10 nM Hg was used in all experiments and Hg was supplied from the same stock (NIST 3133).

For phototrophically-grown cultures, the reactant pool δ^202^Hg^II^ increased steadily over time suggesting cells preferentially reduced lighter Hg^II^ (**Fig 2A**). A mirror trend was observed for δ^202^Hg^0^, which was depleted in ^202^Hg^0^ at the beginning of the experiment, became enriched with heavier isotopes as the reaction proceeded, and eventually approached the initial isotopic composition of the NIST 3133 standard (0‰) (**Fig 2B**).

Curiously, the δ^202^Hg^0^ for f_r_ = 0.94 at 3h in the two phototrophic live cell replicates showed that the product pool was initially enriched with the ^202^Hg^0^ isotope (**Fig 2B**). A similar pattern has been observed in iron reducing bacteria that preferentially reduced heavier Fe^III^ (51) and the model anaerobic Hg methylator and reducer *Geobacter sulfurreducens* PCA (37, 40). The most recent work on *G. sulfurreducens* has shown that uptake of Hg selects for lighter Hg isotopes but cells can also access an isotopically heavier pool of Hg^II^ from the dissolved phase (37). It is possible that a similar process is occurring for phototrophically-grown *H. modesticaldum* wherein cells access an isotopically heavier pool of Hg during equilibrium binding of Hg^II^ to the outside of the cell, followed by an alternative uptake process that selects for lighter pools of Hg. Based on these results, we included all time points in isotopic enrichment calculations for Hg^II^ but omitted the 3h time point in calculations for Hg^0^ given that other cellular fractionation processes could be occurring (**Fig S3, Fig S4**, and **Table S3**).

Positive MDF was also observed for chemotrophic cultures but fractionation patterns differed strongly between replicates. Replicate #1 showed an initial depletion in reactant pool δ^202^Hg^II^ and became progressively enriched with ^202^Hg^II^ relative to the NIST 3133 suggesting lighter isotopes were still preferentially reduced (**Fig 2C**). Note that the second δ^202^Hg^II^ data point available for replicate #1 occurs at f_r_=0.72 due to the rapid rate of Hg^II^ reduction leading to substantially less Hg remaining in the reactor (**Fig 1E** and **Fig 2C**). Results for replicate #2 further suggested that lighter Hg^II^ isotopes were preferentially reduced with δ^202^Hg^II^ slightly increasing at the beginning of the experiment before rising to a maximum of 0.72 ‰ between f_r_ = 0.85 at 12h and 0.53 at 24h (**Fig 2C**).

The trends for δ^202^Hg^0^ in chemotrophic cultures mirrored what was observed for δ^202^Hg^II^. δ^202^Hg^0^ from replicate #1 showed that the product pool was initially enriched with heavy ^202^Hg^0^, further suggesting cells may have accessed a readily bioavailable pool of heavy Hg^II^, before undergoing a pronounced depletion and progressive enrichment with heavy isotopes similar to phototrophic cultures (**Fig 2B, D**). In contrast, the δ^202^Hg^0^ values for cells in replicate #2 suggest that the product pool was depleted in ^202^Hg^0^ before undergoing enrichment for heavier Hg^0^ isotopes reaching a maximum of 1.73 ‰ (**Fig 2D**).

The enrichment in ^202^Hg^0^ exceeded the initial composition of the NIST 3133 standard (0 ‰) suggesting fractionation deviated from the predicted Rayleigh fractionation for Hg reduction (**Table S4**). Recent work with *G. sulfurreducens* has shown that subcellular partitioning processes contribute to Hg isotope fractionation and cannot be detected at the level of THg pool (37). In our work, Hg reduction provided such a strong isotopic shift that we could detect fractionation in the THg pool, however we cannot discount the contributions of subcellular partitioning processes, which are smaller in magnitude (<0.10 ‰). Such processes could be driving the depletion of ^202^Hg^II^ early on for chemotrophic cells in replicate #1 where low cell densities may have amplified uptake and adsorption-driven fractionation (**Fig 1F**). It is more challenging to discuss contributions of such processes for replicate #2, where the low cumulative Hg^0^ production at f_r_ = 0.94 and 0.88 (3h and 6h time points, respectively) precluded our ability to measure Hg^0^ isotope fractionation (**Fig 1E** and **Table S2**). Despite this limitation, the high δ^202^Hg^0^ value obtained for the final timepoint for replicate #2 reinforces that additional processes are contributing to fractionation under chemotrophic conditions that warrant further exploration.

Our experimental results suggest that Hg^0^ oxidation is not a major contributor at earlier stages of reduction (i.e., 3-12 hours) but may be contributing to isotope fractionation at later time points associated with longer Hg residence times in the reactor. Under phototrophic conditions, the measured values of δ^202^Hg^II^ at 48h were isotopically heavier than predicted values, which suggests Hg^0^ oxidation may be occurring (**Table S4**). Under fermentative conditions the potential contribution of Hg oxidation is unclear, since a consistent enrichment was not observed between replicates (**Table S4**). The variability in isotope results for later time points in both replicates suggests processes other than reduction may be occurring.

Although Hg^0^ oxidation has never been observed in anoxygenic phototrophs, it has been observed in other model chemotrophic anaerobes (49). Recent work showed that thiol-bearing molecules preferentially oxidize heavy ^202^Hg^0^, leading to negative MDF, in addition to a MIF signal (52). Although we did not observe any negative MDF or MIF in our experiments (**Table S1** and **Table S2**), we acknowledge that such processes have the potential to contribute to net fractionation alongside internal partitioning related to uptake or adsorption processes. These results are an important first step to resolving the physiological processes that drive these Hg transformations in *H. modesticaldum* and we plan to carry out additional experiments to assess the contributions of uptake, adsorption, and redox transformations to the net Hg isotope fractionation observed in the future.

### Comparing enrichment factors for abiotic and biotic Hg transformations

The average isotopic enrichment factor calculated using the ratio of product to reactant (ε_p/r_) ± 2 standard deviations (2SD) for MDF in the reactant pool Hg^II^ during anoxygenic photosynthesis (ε_AP_) was −0.80 ± 0.56 ‰ whereas the average isotopic enrichment factor obtained during fermentation (ε_FM_)was −1.13 ± 0.50 ‰ (**Fig 3, Fig S3, Table S3**, and **Table S5**).

**Figure 3:**
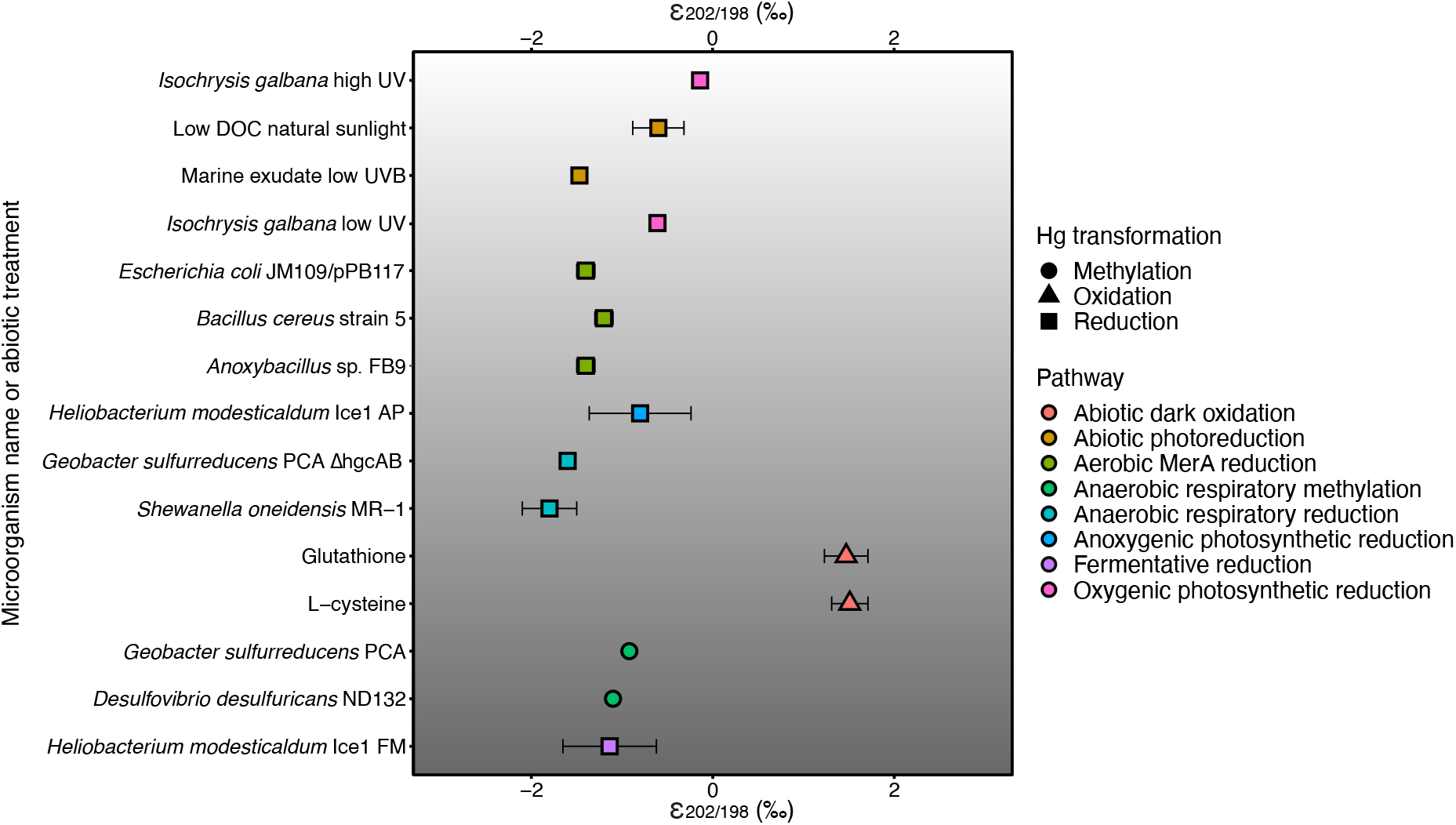
Compilation of isotopic enrichment factors (ε) for abiotic and biotic Hg transformations. Hg transformations are denoted by different shapes and specific metabolic and chemical pathways have been colour coded. Error bars denote 2 standard deviations for studies where this data was available. In the event that data was presented as a standard error, standard deviation was calculated based on the sample size indicated in each study. Abbreviations on the y-axis denote the following: UV (ultraviolet light), DOC (dissolved organic carbon), AP (anoxygenic photosynthesis), ΔhgcAB (no *hgcAB* gene cluster), and FM (fermentation). The grey-shading in this figure represents light and oxygen. Lighter regions represent habitats where light and oxygen are present and darker regions represent where they are not. The values in this figure were obtained from the following studies: abiotic dark oxidation (52); abiotic photoreduction (32, 41); aerobic MerA reduction (34, 36); anaerobic respiratory methylation (40); anaerobic respiratory reduction (36, 37); anoxygenic photosynthetic reduction (this study); fermentative reduction (this study); oxygenic photosynthetic reduction (41). The raw data in this figure are in **Table S5**.

The enrichment factor could not be calculated using the product Hg^0^ pool for fermentative cultures and overestimated the value in phototrophic cultures (−2.86 ± 1.32 ‰) (**Fig S4** and **Table S3**). Previous work has shown that using the product pool can overestimate the fractionation factor due to difficulty obtaining analytical replicates of gaseous Hg^0^ and the complexity of quantitatively collecting this pool (53). As such, we carried out comparisons for enrichment factors using the reactant Hg^II^ pool. Ideally, we would have been able to obtain more replicates of our experiments but were unable to do so due the challenges stated previously. It is important to note that low numbers of replicates are routinely discussed in isotope geochemistry and the variability from our results is in line with previous work (**Fig 3** and **Table S5**).

Our initial motivation for this study was to test if phototrophic and fermentative Hg reduction led to markedly different isotope enrichment factors and establish whether different underlying processes supported different metabolic Hg reduction pathways. When comparing ε values, which are used as an index to distinguish between different isotope fractionation processes, we report a difference of 0.33 ‰ between ε_AP_ and ε_FM_ (**Fig 3** and **Table S5**). This small difference in enrichment factors supports that common processes are controlling Hg reduction during anoxygenic photosynthesis and fermentation in *H. modesticaldum*.

In our previous work with *H. modesticaldum* we demonstrated that inhibiting pyruvate ferredoxin oxidoreductase, the enzyme responsible for the production of reduced ferredoxin, greatly hampered Hg^II^ reduction during anoxygenic photosynthesis and fermentation (23). It is possible that the similarity between ε_AP_ and ε_FM_ is the result of reactions originally involving the same redox cofactors (e.g. ferredoxins) but with Hg^II^ being reduced by different electron donors specific to each metabolism. Such reactions could involve direct electron transfer from ferredoxin to Hg^II^ or from enzymes directly or indirectly relying on ferredoxin (23, 54). It is also possible that the net isotopic enrichment stems from a combination of multiple processes involved in Hg transport (e.g. adsorption, uptake, and efflux) that would occur regardless of the intracellular transformations in question. Our results suggest that the relative contributions of these processes may be more pronounced during fermentation compared to anoxygenic photosynthesis although more controlled experiments are required to constrain the variability in the isotope fractionation observed.

At the broader scale, the isotopic enrichment observed for both metabolisms tested in *H. modesticaldum* falls between the values observed for abiotic and microbial Hg reduction pathways (**Fig 3** and **Table S5**). The isotopic enrichment observed for phototrophic *H. modesticaldum* is slightly lower than photoreduction in the presence of dissolved organic carbon (−0.80 ± 0.56 ‰ vs −0.60 ± 0.20 ‰, respectively) (32) but higher than photoreduction in the presence of marine exudates and UVB light (−0.80 ± 0.56 ‰ vs −1.47 ‰, respectively) (41) (**Fig 3** and **Table S5**). When comparing Hg reduction in the anoxygenic phototroph *H. modesticaldum* to the oxygenic phototrophic green alga *Isochrysis galbana*, the isotopic enrichment observed is also slightly lower (−0.80 ± 0.56 ‰ vs −0.14 to −0.60 ‰, respectively) (41) (**Fig 3** and **Table S5**). Though this similarity suggests anoxygenic and oxygenic photosynthetic Hg reduction share a common physiological pathway, the presence of MIF in *I. galbana* due to free radical generation within the cell rules out this possibility (41). (**Table S1**). Thus, Hg reduction during anoxygenic photosynthetic Hg reduction could be distinguished from other photo-induced reduction pathways due to the lack of MIF tracer.

When compared with previous work on microbial Hg reduction in pure cultures, the isotopic enrichment observed for phototrophically-grown *H. modesticaldum* was higher than what has previously been reported for aerobic *mer*-mediated reduction and anaerobic respiratory Hg reduction (−0.80 ± 0.56 ‰ vs −1.80 to −1.20 ‰, respectively) (34, 36, 37) (**Fig 3** and **Table S5**). Interestingly, the isotopic enrichment observed in fermentatively-grown *H. modesticaldum* (−1.13 ± 0.50 ‰) was closer to aerobic *mer*-mediated reduction (−1.40 to −1.20 ‰) (34) than to anaerobic respiratory Hg reduction in *Shewanella oneidensis* MR-1 and a modified strain of *G. sulfurreducens* PCA incapable of Hg methylation (−1.80 to −1.60 ‰) (**Fig 3** and **Table S5**) (36, 37). An additional line of evidence supporting that Hg^0^ oxidation is unlikely to be contributing to the isotope signature observed in our system are the ε values for thiol mediated Hg^0^ oxidation in the dark. These values are considerably larger than what we observed for *H. modesticaldum* (1.51 ± 0.20 ‰ for cysteine and 1.47 ± 0.24 ‰ for glutathione) (**Fig 3** and **Table S5**).

These comparisons illustrate that isotopic enrichment for microbial Hg reduction pathways that only display MDF fall within a narrow range (i.e. −0.80 to −1.80 ‰) (**Fig 3** and **Table S5**). We find this to be a strikingly small difference given the ecological diversity of the model organisms and variety of growth conditions used to study microbial Hg reduction. The similar ε values reported for fermentation in this study and the *mer* operon are also noteworthy as they suggest nearly identical isotope enrichment can occur in aerobes harbouring dedicated enzymatic machinery to reduce Hg and anaerobes where *mer*-based strategies are largely absent.

Our study also shows that similar isotopic enrichment can be observed for different anaerobic Hg transformations. The anaerobic Hg methylating chemotrophs *G. sulfurreducens* PCA and *Desulfovibrio desulfuricans* ND132 displayed ε signatures (i.e. −0.92 and −1.1‰, respectively) that fall within the range of ε values we report in our study (i.e. −0.80 to −1.13 ‰) (**Fig 3** and **Table S5**) (40). The similarity in enrichment signatures suggests that common processes may affect the delivery of inorganic Hg substrates to intracellular sites independently of the Hg transformations in question. Although it is outside of the scope of the current study, identifying the nature of these processes may provide insights into similar controls that exist for Hg uptake and transformation in anaerobes.

Our work showcases the strengths but also the challenges of using isotope fractionation-based approaches to decipher cryptic Hg cycling pathways. To illustrate the challenges of using isotopes to track Hg cycling pathways in the environment, we visualized our comparison of ε values in the context of a vertically redox stratified habitat (e.g. a water column) (**Fig 3**). Through Hg stable isotope fractionation, we can distinguish between abiotic and biotic redox reaction pathways that control Hg’s availability in methylation hotspots, such as the limit of the photic zone, on the bases of MIF signatures associated with photochemical reactions. In contrast, teasing apart biological pathways that contribute to the net isotope signature of Hg^II^ in the environment is more challenging because competing MDF processes occur in both dark and photo-mediated pathways where Hg^II^ is a reactant (e.g. reduction and methylation) and a product (e.g. demethylation and oxidation). This overlap makes it difficult to translate δ^202^Hg values observed under controlled conditions that are sensitive to changes in cellular physiology to environments where pools of bioavailable Hg are not homogenous due to variable mixing.

## CONCLUSION

In this study we have provided the first evidence that common physiological processes drive MDF during Hg reduction associated with anoxygenic photosynthesis and fermentation in the model anaerobe *H. modesticaldum*. We show that isotopic enrichment during anoxygenic phototrophic Hg reduction leads to an intermediate fractionation process that lies between photochemical reactions and dark microbial Hg reduction. We also show that fermentative Hg reduction leads to isotopic enrichment that is similar to aerobic *mer*-mediated reduction.

Although current analytical capacities to measure stable isotope fractionation make applying such approaches to the environment challenging, this is a burgeoning field that has advanced considerably in terms of sensitivity and the different Hg species that can be targeted in the last 10 years. Isotope fractionation approaches have proved extremely valuable for teasing apart the complex interwoven processes that drive microbe-mediated Hg transformations and as seen in our study, highlighting common processes that control Hg bioavailability to microbes.

In future work, it will be important to isolate the different steps involved in Hg uptake and transformation by rigorously controlling Hg availability and cellular physiology. Leveraging molecular tools alongside isotopes will be a useful strategy to address these knowledge gaps and the recent availability of genetically tractable *H. modesticaldum* deletion mutants offers a promising means to do so (55). In that regard, gene deletion approaches targeting redox-active enzymes could be used to characterize the physiological components contributing directly to Hg^II^ reduction, whereas targeting homologues for proteins involved in Hg uptake in other model anaerobes could be the key to identifying entry routes for Hg into *H. modesticaldum* (56, 57).

In addition to gene deletions, whole-genome, transcriptome, and proteome approaches could be combined with isotopic analyses in pure cultures to hone in on potential biomarkers for Hg transformation pathways that remain poorly understood. Such biomarkers could be used to examine publicly available genomic datasets to assess the distribution of anaerobic Hg reducers in modern day environments and rock records, the latter serving as an archive for Hg isotope signatures and the microbial community involved in historical Hg cycling (31, 58). These ancient environments were likely dominated by anoxygenic phototrophs that were important primary producers before the rise of oxygen on Earth. By using isotopes to create a window into the past, we can compare the selective forces that gave rise to fortuitous Hg reduction pathways tied to anaerobic metabolisms and dedicated detoxification strategies such as the *mer* operon.

## ACKNOWLEDGMENTS

Our work was funded by NSERC Discovery and Accelerator grants, CFI funding to AJP and NSERC graduate scholarship to DSG. Instrumentation and lab operations for Hg isotope analyses were supported by the USGS Toxic Substances Hydrology Program. Any use of trade, firm, or product names is for descriptive purposes only and does not imply endorsement by the U.S. Government.

## AUTHOR CONTRIBUTIONS

DSG, AJP, SEJ and MTT designed all experiments; DSG and NCL carried out bioreactor experiments and data analyses; SEJ and MTT carried out total Hg and stable isotope analyses; DSG, AJP, SEJ and MTT wrote the manuscript.

## SUPPORTING INFORMATION CAPTIONS

**Figure S1:** Schematic of the bioreactor setup used to measure Hg stable isotope fractionation for Hg^II^ and Hg^0^ during the anaerobic reduction of Hg^II^.

**Figure S2**: **Total Hg mass balance for isotope fractionation experiments under phototrophic and chemotrophic growth conditions**. The average recovery across all experiments was 94.40 ± 9.50 % (n=7) and is shown by the dashed red line.

**Figure S3: Mass dependent fractionation of ^202^Hg^II^ in phototrophically (A, B) and chemotrophically (C, D) grown *H. modesticaldum***. δ^202^Hg values for Hg^II^ are plotted as open system Rayleigh fractionation models with ln(R/R_0_) vs ln(f_r_). The amount of Hg^II^ remaining in the reactor was used to calculate f_r_ (see Methods). The 0h time point for chemotrophic live cells replicate #1 was omitted from the regression fitted to the data for Hg^II^ (C) as this suggested an alternative process was contributing to isotopic fractionation earlier on in the experiment.

**Figure S4: Mass dependent fractionation of ^202^Hg^0^ in phototrophically (A, B) and chemotrophically (C, D) grown *H. modesticaldum***. δ^202^Hg values for Hg^0^ are plotted as open system Rayleigh fractionation models with ln(R/R_0_) vs ln(f_r_). The amount of Hg^II^ remaining in the reactor was used to calculate f_r_ (see Methods). The 3h time point for phototrophic cells in replicates #1 and 2 were omitted from the regression fitted to the data for Hg^0^ (A, B) as this suggested an alternative process was contributing to isotopic fractionation earlier on in the experiment. No significant regressions were obtained for chemotrophic cells and as such no models are presented.

**Table S1**: Total Hg concentrations and Hg isotope ratios for the reactant pool Hg^II^ in phototrophic and chemotrophic growth conditions experiments. Each line corresponds to an individual sample and technical duplicates were taken for each time point. Std represents the standard deviation from analytical replicates on the MC-ICP-MS. Abbreviations denote Metabo. = Metabolism; Photo. = Phototrophy; Chemo. = Chemotrophy; Treat. = Treatment; Hg conc. = Hg concentration, rep = replicate; Avg = average; Std = Standard deviation; ctl = control, MED = medium. δ^202^Hg values in **bold** denote those that deviate substantially from the composition of the NIST 3133 standard in medium spiked controls.

**Table S2:** Total Hg concentrations and Hg isotope ratios for the Hg^0^ pool in phototrophic and chemotrophic growth conditions experiments. Each line corresponds to the Hg present on two gold traps used to sample each time point in parallel. Std represents the standard deviation from analytical replicates on the MC-ICP-MS. Abbreviations denote the following: Metabo. = Metabolism; Photo. = Phototrophy; Chemo. = Chemotrophy; Treat. = Treatment; Hg conc. = Hg concentration, rep = replicate; Avg = average; Std = Standard deviation.

**Table S3:** Summary of regression models for mass dependent fractionation of ^202^Hg^II^ and ^202^Hg^0^ in phototrophically and chemotrophically grown *H. modesticaldum* cultures. P-values for regressions that displayed a significant (p<0.05) fit are shown in **bold**.

**Table S4:** Comparison of measured and calculated isotope fractionation values for the reactant pool of Hg^II^ using a mass balance approach. Measured values of Hg concentration and δ^202^Hg are represented as an average of replicates from experiments presented in Tables S1 and S3. Predicted values for media were obtained using the equations outlined for mass balance calculations. Differences between the measured and predicted values that are substantially greater than the analytical variability of measurements (2SD = 0.08) are shown in **bold**. Abbreviations denote Photo. = phototrophic, Chemo. = chemotrophic, rep = replicates, Avg. = average, Pred. = predicted.

**Table S5:** Comparison of enrichment factors and measures of variability used in this study and the original cited work. Values in bold represent calculation ratios that differ between the original article cited and the current study. Abbreviations: SD = standard deviation, SE = standard error, p = product, r = reactant, Ox. = oxygenic, Anox. = anoxygenic, PS = photosynthesis.

## Notes

### Competing Interest Statement

The authors have declared no competing interest.

