## Supporting Information for "Common physiological processes control mercury reduction during photosynthesis and fermentation"

**SUPPORTING METHODS**

Bioreactor setup and control samples for isotope fractionation

MC-ICP-MS analytical expectations

Isotope fractionation factor calculations

Isotope mass balance calculations

**SUPPORTING FIGURES AND TABLES**

**Figure S1**: Schematic of bioreactor setup

**Figure S2**: Total Hg mass balance for isotope fractionation experiments under phototrophic and chemotrophic growth conditions

**Figure S3**: Regression models used to calculate isotopic enrichments during mass dependent fractionation of $^{202}Hg^{II}$ in phototrophically and chemotrophically grown *H. modesticaldum* cultures

**Figure S4**: Regression models used to calculate isotopic enrichments during mass dependent fractionation of $^{202}Hg^{0}$ in phototrophically and chemotrophically grown *H. modesticaldum* cultures

**Table S1**: Total Hg concentrations and Hg isotope ratios for the reactant pool $Hg^{II}$ in phototrophically and chemotrophically grown cultures of *H. modesticaldum*

**Table S2**: Total Hg and Hg isotope ratios recovered on gold traps for the product pool $Hg^{0}$ in phototrophically and chemotrophically grown cultures of *H. modesticaldum*

**Table S3**: Summary of regression models for mass dependent fractionation of $^{202}Hg^{II}$ and $^{202}Hg^{0}$ in phototrophically and chemotrophically grown *H. modesticaldum* cultures

**Table S4:** Isotope mass balance to test for deviations from Rayleigh fractionation model

**Table S5:** Compilation of isotope enrichment factors and measures of variability

### SUPPORTING METHODS

#### Bioreactor setup and control samples for isotope fractionation

In contrast to our previous work capturing pM levels of $Hg^0$ in real-time using bioreactors, capturing nM levels of $Hg^0$ for isotope analyses required the use of gold traps. Our preliminary efforts showed that single gold traps were insufficient to study isotope fractionation because high levels of $Hg^0$ produced early on in experiments could break through the trap due to saturation and high $N_2$ purging rates. As such, two gold traps were employed to capture all $Hg^0$ (**Fig S1**). When setting up experiments, bioreactors were purged continuously for 2h to remove background $Hg^0$ prior to inoculation after which a subsample was taken to measure the background total Hg (THg) that was present in the medium. Afterwards, the reactor was purged for 1h to remove any background $Hg^0$ that may have infiltrated the experimental setup due to subsampling. To control for possible fractionation due to medium composition, an additional subsample of growth medium was spiked directly with the National Institute of Standards and Technology (NIST) 3133 standard (initial Hg concentration of 23.28 μM) to a final concentration 10 nM Hg. The isotopic compositions of these samples were compared to high purity water spiked with the same NIST 3133 standard to verify that no isotope fractionation occurred as a result of the growth conditions employed (**Table S1**).

**MC-ICP-MS analytical expectations**

The instrument was tuned (i.e. sample gases, stage positions, and lenses) to achieve the optimal $^{202}Hg$ voltage of approximately 1V for a 5 nM Hg solution. A secondary standard (NIST RM 8610) was measured within each sample batch to ensure instrument stability and accuracy. Average values obtained for NIST 8610 with 2 standard deviations were ($\delta^{202}Hg$= -0.53 ± 0.08; $\Delta^{199}Hg$ =-0.02 ± 0.06; $\Delta^{200}Hg$ =0.00 ± 0.04; $\Delta^{201}Hg$ =-0.04 ± 0.06; $\Delta^{204}Hg$ =0.00 ± 0.14, 2SD, n = 35), which agreed with certified values. Additionally, desorption check standards (NIST 3133) for $Hg^0$ traps were measured to verify that no fractionation occurred during gold trap processing. Standards displayed average values in line with their initial isotopic composition ($\delta^{202}Hg$= -0.04 ± 0.10; $\Delta^{199}Hg$ =0.03 ± 0.06; $\Delta^{200}Hg$ =0.0 ± 0.06; $\Delta^{201}Hg$ =0.01 ± 0.06; $\Delta^{204}Hg$ =-0.04 ± 0.14, 2SD, n = 15) indicating no fractionation during processing.

In some cases, medium controls spiked directly with 10 nM of the NIST 3133 standard showed positive $\delta^{202}Hg$ values that ranged from 0.5 to 0.8 ‰, suggesting isotope fractionation was occurring due to reactions withing the growth medium or sampling vessel (**bold values** in **Table S1**). This fractionation seems to be an artefact associated with the preparation of those specific samples as it did not occur consistently in all medium controls made from the same recipe and no fractionation was observed for any of the water-controls treated in the same fashion (**Table S1**). Furthermore, this fractionation seems to be absent from all 0h time points taken from the bioreactor (**Table S1**). Although we cannot explain why this fractionation occurred, the absence of consistent fractionation in the medium blanks and 0h time points in our experiments means we can still compare isotope fractionation from the bioreactor with confidence.

**Isotope fractionation factor calculations**

Fractionation factors for Hg reduction were calculated using the following equations for the reactants and products, respectively:

$$\ln(R_{media}/R_0) = \ln(f_r) * (1/\alpha_{r/p} - 1) \quad \textbf{(Eq S1)}$$

$$\ln(R_{gas}/R_0) = \ln(f_r) * (1/\alpha_{r/p} - 1) + \ln(1/\alpha_{r/p}) \quad \textbf{(Eq S2)}$$

Where $R_{media}$ is the relative isotope ratio of $Hg^{II}$ in the media at a given time point, $R_{gas}$ is the relative isotope ratio of $Hg^0$ collected onto gold traps at a given time point, $R_0$ is the initial relative isotope ratio in the reactor at the start of the experiment, $f_r$ is the fraction of $Hg^{II}$ remaining in the reactor, and $\alpha_{r/p}$ is the kinetic fractionation factor for $(^{202}Hg/^{198}Hg)_{reactant}/(^{202}Hg/^{198}Hg)_{product}$. The $\alpha_{r/p}$ value can be converted to $\varepsilon_{p/r}$, which is the isotope enrichment factor for the product pool, using:

$$\varepsilon_{p/r} = [(\alpha_{r/p}-1)*1000]/-1 \quad \textbf{(Eq S3)}$$

It was noted that the slope of the regression was equivalent to the $\varepsilon_{p/r}$ when multiplied by 1000 to convert to per mille (‰) notation (**Eq 5** in the main text).

Our review of the literature showed that several studies present the ratio of reactant to product rather than product to reactant for ε, the latter being the method used in this study. Furthermore, our review showed that isotopes studies published to date have differing degrees of replications and employ different measures of variability. We have attempted to convert all ε values from the literature to values that represent the conversion of reactants to products according to previous calculations used for microbial $Hg^{II}$ reduction (1, 2) (see **Table S5**).

If the isotope fractionation factor ($\alpha_{202/198}$) was presented instead of ε, α was converted to ε based on **Eq S3.** If the ratio of product to reactants was employed, the reciprocal value of α was used instead. Where sample size was available, we also attempted to convert all measures of standard error to measures of 2 x standard deviation, as is the convention set by previous work comparing different pathways for microbial Hg transformation (1, 2).

**Mass balance calculations**

Mass balance calculations were used to predict the isotopic composition of the $Hg^{II}$ pool to determine if Hg reduction was the dominant reaction pathway during phototrophic and chemotrophic experiments. For the calculations it was assumed that the sum of isotopic products would be equivalent to the reactants:

$\delta^{202}Hg_R(f_R) + \delta^{202}Hg_P(f_P) = 0$ (**Eq S4**)

Where R represents the reactant pool of $Hg^{II}$ in the reactor and P represents the product pool of $Hg^0$ for the isotopic composition ($\delta^{202}Hg$) and fraction remaining (f) of each pool. This equation was simplified to solve for the predicted isotopic composition of $\delta^{202}Hg_R$ using the values presented in **Table S5**:

$\delta^{202}Hg_R = -[\delta^{202}Hg_P(f_P)]/ f_R$ (**Eq S5**)

If only Hg reduction was driving the isotopic fractionation within these cultures the measured and predicted values of $\delta^{202}Hg_R$ for $Hg^{II}$ in the media should be similar. Large differences between the calculated and measured values would indicate that other reactions may be causing the calculations to under or overestimate the $Hg^{II}$ pool and these values are highlighted in **bold** in **Table S4**.

### SUPPORTING FIGURES

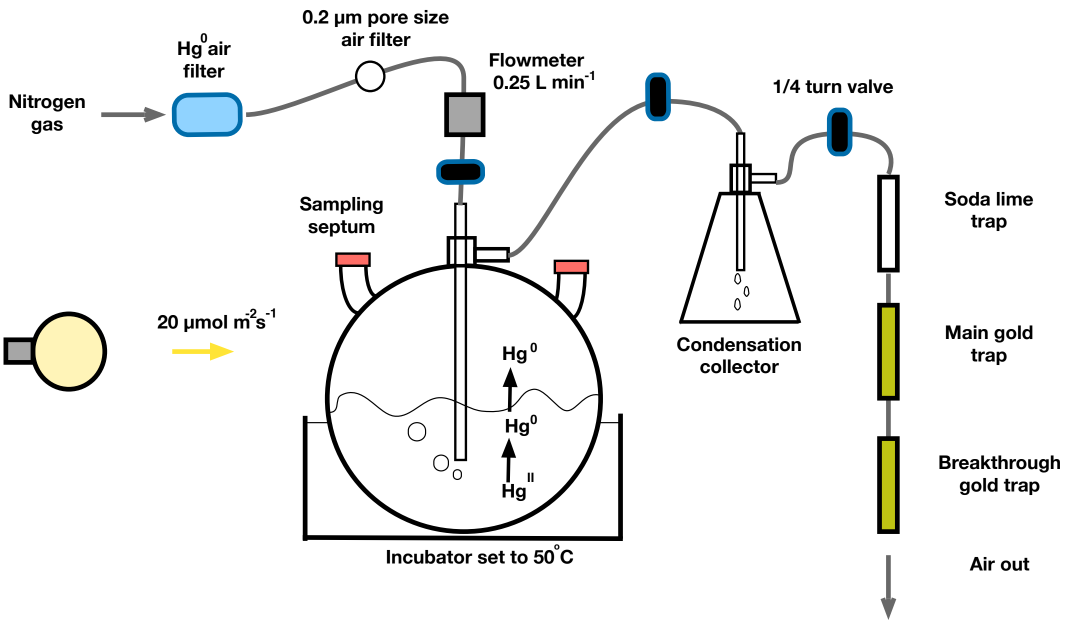


**Figure S1:** Schematic of the bioreactor setup used to measure Hg stable isotope fractionation for $Hg^{II}$ and $Hg^0$ during the anaerobic reduction of $Hg^{II}$.

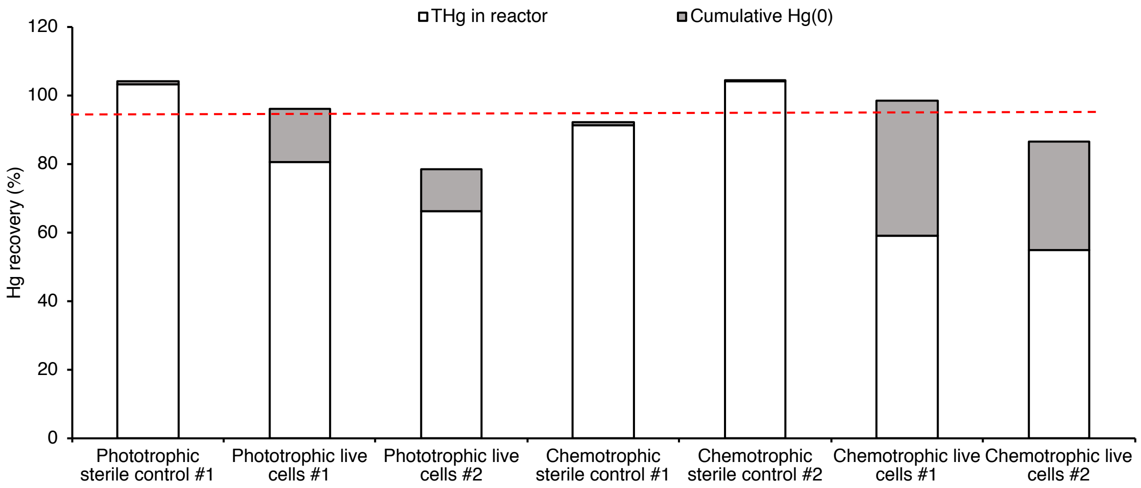


**Figure S2**: **Total Hg mass balance for isotope fractionation experiments under phototrophic and chemotrophic growth conditions.** The average recovery across all experiments was 94.40 ± 9.50 % (n=7) and is shown by the dashed red line.

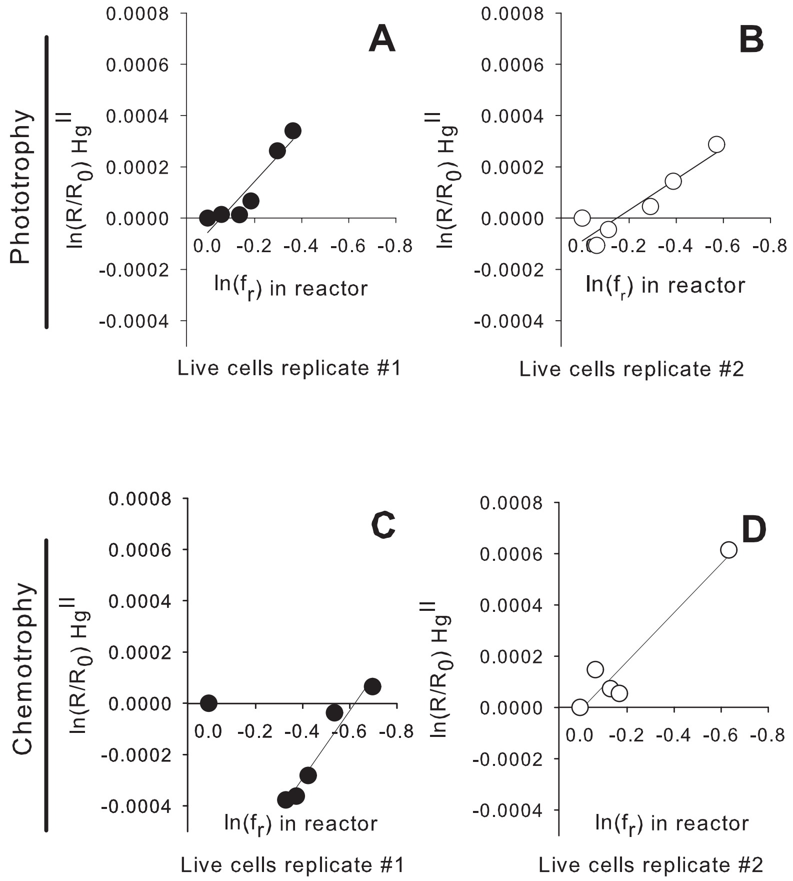


**Figure S3: Mass dependent fractionation of $^{202}Hg^{II}$ in phototrophically (A, B) and chemotrophically (C, D) grown *H. modesticaldum*.** $\delta^{202}Hg$ values for $Hg^{II}$ are plotted as open system Rayleigh fractionation models with $\ln(R/R_0)$ vs $\ln(f_r)$. The amount of $Hg^{II}$ remaining in the reactor was used to calculate $f_r$ (see Methods). The 0h time point for chemotrophic live cells replicate #1 was omitted from the regression fitted to the data for $Hg^{II}$ (C) as this suggested an alternative process was contributing to isotopic fractionation earlier on in the experiment.

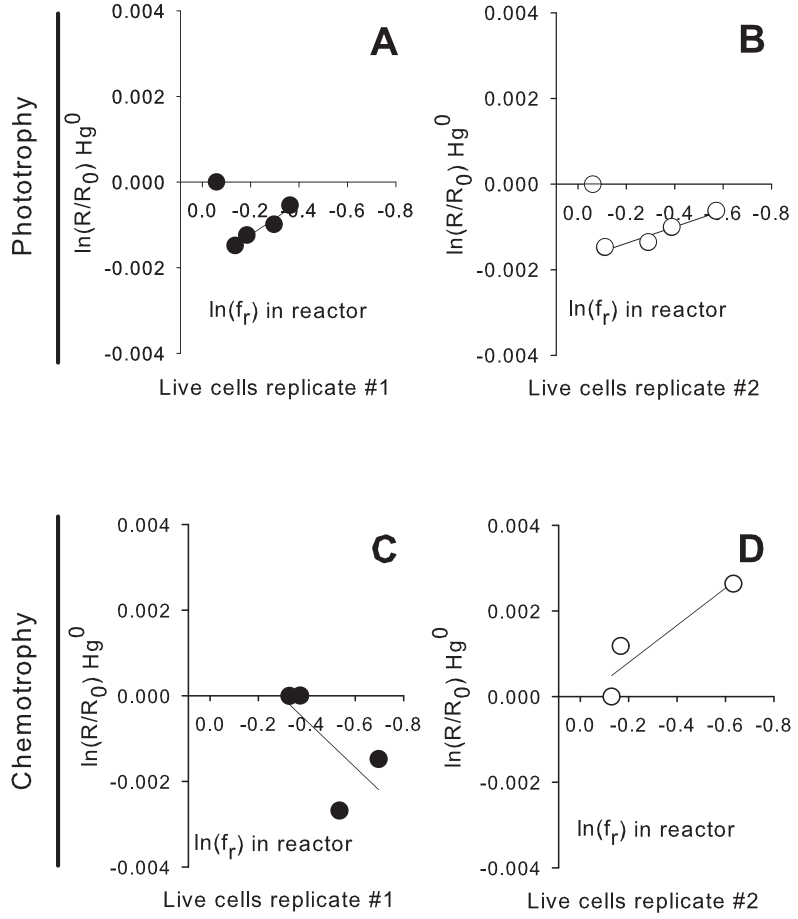


**Figure S4: Mass dependent fractionation of $^{202}Hg^0$ in phototrophically (A, B) and chemotrophically (C, D) grown *H. modesticaldum*.** $\delta^{202}Hg$ values for $Hg^0$ are plotted as open system Rayleigh fractionation models with $\ln(R/R_0)$ vs $\ln(f_r)$. The amount of $Hg^{II}$ remaining in the reactor was used to calculate $f_r$ (see Methods). The 3h time point for phototrophic cells in replicates #1 and 2 were omitted from the regression fitted to the data for $Hg^0$ (A, B) as this suggested an alternative process was contributing to isotopic fractionation earlier on in the experiment. No significant regressions were obtained for chemotrophic cells and as such no models are presented.

**Table S1**: Total Hg concentrations and Hg isotope ratios for the reactant pool $Hg^{II}$ in phototrophic and chemotrophic growth conditions experiments. Each line corresponds to an individual sample and technical duplicates were taken for each time point. Std represents the standard deviation from analytical replicates on the MC-ICP-MS. Abbreviations denote Metabo. = Metabolism; Photo. = Phototrophy; Chemo. = Chemotrophy; Treat. = Treatment; Hg conc. = Hg concentration, rep = replicate; Avg = average; Std = Standard deviation; ctl = control, MED = medium. $\delta^{202}$Hg values in **bold** denote those that deviate substantially from the composition of the NIST 3133 standard in medium spiked controls.

| Metabo. | Treat. | Sample | Hg conc. (nM) | $\delta^{202}$Hg Avg | $\delta^{202}$Hg Std | $\Delta^{199}$Hg Avg | $\Delta^{199}$Hg Std | $\Delta^{200}$Hg Avg | $\Delta^{200}$Hg Std | $\Delta^{201}$Hg Avg | $\Delta^{201}$Hg Std | $\Delta^{204}$Hg Avg | $\Delta^{204}$Hg Std |
|---|---|---|---|---|---|---|---|---|---|---|---|---|---|
| Photo. | Sterile ctl | MED. ctl | 9.01 | **0.8329** | 0.0580 | -0.0453 | 0.0317 | -0.0125 | 0.0112 | -0.0325 | 0.0277 | 0.3219 | 0.0420 |
| | | H2O ctl | 10.43 | -0.0380 | 0.0474 | -0.0463 | 0.0268 | 0.0093 | 0.0123 | -0.0389 | 0.0296 | 0.1485 | 0.0578 |
| | | TIME 0HR | 8.82 | -0.0291 | 0.0441 | -0.0024 | 0.0270 | -0.0033 | 0.0289 | 0.0013 | 0.0179 | 0.0426 | 0.0396 |
| | | TIME 0HR | 9.03 | 0.0201 | 0.0504 | -0.0626 | 0.0302 | -0.0236 | 0.0122 | -0.0176 | 0.0183 | 0.1059 | 0.0399 |
| | | TIME 3HR | 9.02 | -0.0283 | 0.0336 | -0.0324 | 0.0425 | -0.0327 | 0.0249 | 0.0047 | 0.0155 | -0.0545 | 0.0583 |
| | | TIME 3HR | 9.50 | 0.0784 | 0.0395 | 0.0757 | 0.0398 | 0.0226 | 0.0192 | 0.0582 | 0.0102 | 0.0249 | 0.0580 |
| | | TIME 6HR | 9.11 | 0.0462 | 0.0376 | 0.0225 | 0.0397 | 0.0191 | 0.0379 | 0.0625 | 0.0258 | 0.0215 | 0.0535 |
| | | TIME 6HR | 8.75 | 0.0339 | 0.0235 | 0.0141 | 0.0180 | -0.0216 | 0.0364 | -0.0056 | 0.0237 | -0.0995 | 0.0519 |
| | | TIME 12HR | 8.97 | 0.0033 | 0.0450 | -0.0121 | 0.0261 | -0.0179 | 0.0379 | -0.0051 | 0.0498 | -0.0183 | 0.0438 |
| | | TIME 12HR | 8.67 | -0.0292 | 0.0295 | 0.0696 | 0.0215 | -0.0053 | 0.0364 | 0.0289 | 0.0496 | -0.0532 | 0.0312 |
| | | TIME 24HR | 9.19 | 0.0441 | 0.0159 | 0.0277 | 0.0244 | 0.0559 | 0.0422 | -0.0023 | 0.0425 | 0.1206 | 0.0487 |
| | | TIME 24HR | 9.15 | -0.0671 | 0.0470 | 0.0241 | 0.0336 | -0.0339 | 0.0192 | -0.0095 | 0.0206 | 0.0894 | 0.0528 |
| | | TIME 48HR | 9.00 | 0.0489 | 0.0303 | 0.1298 | 0.0307 | 0.0182 | 0.0268 | -0.0084 | 0.0228 | 0.0262 | 0.0537 |
| | | TIME 48HR | 8.89 | 0.0559 | 0.0295 | 0.0412 | 0.0364 | 0.0395 | 0.0139 | 0.0182 | 0.0169 | 0.0776 | 0.0650 |
| | Live cells rep #1 | MED. ctl | 9.76 | -0.0175 | 0.0279 | -0.0106 | 0.0658 | -0.0136 | 0.0338 | 0.0193 | 0.0187 | 0.0654 | 0.0170 |
| | | H2O ctl | 10.28 | 0.0900 | 0.0277 | 0.0567 | 0.0610 | 0.0701 | 0.0350 | 0.0627 | 0.0183 | -0.0207 | 0.0510 |
| | | TIME 0HR | 9.48 | 0.0424 | 0.0391 | -0.0064 | 0.0560 | -0.0394 | 0.0178 | -0.0387 | 0.0099 | 0.0022 | 0.0130 |
| | | TIME 0HR | 9.50 | 0.0197 | 0.0272 | -0.0190 | 0.0308 | 0.0263 | 0.0151 | 0.0206 | 0.0090 | -0.0373 | 0.0425 |
| | | TIME 3HR | 8.71 | 0.0831 | 0.0618 | 0.0219 | 0.0157 | 0.0280 | 0.0090 | -0.0258 | 0.0061 | -0.0080 | 0.0634 |
| | | TIME 3HR | 9.20 | 0.0073 | 0.0533 | 0.0068 | 0.0276 | -0.0188 | 0.0231 | -0.0407 | 0.0462 | -0.0234 | 0.0652 |
| | | TIME 6HR | 8.03 | 0.0629 | 0.0336 | -0.0061 | 0.0294 | 0.0029 | 0.0398 | 0.0150 | 0.0443 | -0.0721 | 0.0751 |
| | | TIME 6HR | 8.48 | 0.0252 | 0.0121 | -0.0437 | 0.0232 | -0.0284 | 0.0399 | 0.0201 | 0.0432 | -0.0494 | 0.0527 |
| | | TIME 12HR | 7.96 | 0.0248 | 0.0464 | -0.0220 | 0.0378 | 0.0046 | 0.0168 | 0.0037 | 0.0433 | -0.0053 | 0.0642 |
| | | TIME 12HR | 7.67 | 0.1710 | 0.0228 | 0.0240 | 0.0279 | 0.0008 | 0.0245 | 0.0023 | 0.0477 | -0.1237 | 0.0316 |
| | | TIME 24HR | 6.87 | 0.2817 | 0.0279 | -0.0238 | 0.0352 | 0.0167 | 0.0370 | -0.0237 | 0.0336 | 0.0320 | 0.0504 |
| | | TIME 24HR | 6.77 | 0.3052 | 0.0244 | -0.0361 | 0.0314 | -0.0062 | 0.0199 | -0.0183 | 0.0405 | -0.0949 | 0.0602 |
| | | TIME 48HR | 6.18 | 0.4454 | 0.0550 | -0.0417 | 0.0454 | 0.0256 | 0.0208 | -0.0276 | 0.0358 | 0.0520 | 0.0595 |
| | | TIME 48HR | 6.34 | 0.2975 | 0.0651 | -0.0981 | 0.0510 | -0.0500 | 0.0230 | -0.0372 | 0.0365 | -0.0202 | 0.0630 |

Continued on next page

**Table S1 (continued)**: Total Hg concentrations and Hg isotope ratios for the reactant pool $Hg^{II}$ in phototrophic and chemotrophic growth conditions experiments. Each line corresponds to an individual sample and technical duplicates were taken for each time point. Std represents the standard deviation from analytical replicates on the MC-ICP-MS. Abbreviations denote Metabo. = Metabolism; Photo. = Phototrophy; Chemo. = Chemotrophy; Treat. = Treatment; Hg conc. = Hg concentration, rep = replicate; Avg = average; Std = Standard deviation; ctl = control, MED = medium. $\delta^{202}$Hg values in **bold** denote those that deviate substantially from the composition of the NIST 3133 standard in medium spiked controls.

| Metabo. | Treat. | Sample | Hg conc. (nM) | $\delta^{202}$Hg Avg | $\delta^{202}$Hg Std | $\Delta^{199}$Hg Avg | $\Delta^{199}$Hg Std | $\Delta^{200}$Hg Avg | $\Delta^{200}$Hg Std | $\Delta^{201}$Hg Avg | $\Delta^{201}$Hg Std | $\Delta^{204}$Hg Avg | $\Delta^{204}$Hg Std |
|---|---|---|---|---|---|---|---|---|---|---|---|---|---|
| Photo. | Live cells rep #2 | MED. ctl | 9.11 | 0.1050 | 0.0521 | 0.0300 | 0.0584 | -0.0316 | 0.0082 | 0.0037 | 0.0298 | -0.0627 | 0.0377 |
| | | H2O ctl | 10.27 | 0.0156 | 0.0113 | -0.0487 | 0.0195 | -0.0521 | 0.0131 | -0.0354 | 0.0489 | -0.0213 | 0.0326 |
| | | TIME 0HR | 9.25 | 0.1193 | 0.0408 | 0.0796 | 0.0494 | 0.0176 | 0.0248 | 0.0030 | 0.0220 | -0.0809 | 0.0452 |
| | | TIME 0HR | 8.89 | 0.0519 | 0.0441 | -0.0582 | 0.0175 | -0.0504 | 0.0243 | -0.0355 | 0.0575 | -0.0003 | 0.0333 |
| | | TIME 3HR | 8.48 | 0.0000 | 0.0646 | 0.0016 | 0.0486 | 0.0121 | 0.0322 | -0.0604 | 0.0577 | -0.1155 | 0.0551 |
| | | TIME 3HR | 8.58 | -0.0428 | 0.0760 | -0.0321 | 0.0504 | 0.0174 | 0.0322 | 0.0322 | 0.0638 | 0.0205 | 0.0638 |
| | | TIME 6HR | 8.35 | 0.0581 | 0.0960 | 0.1027 | 0.0459 | 0.0074 | 0.0199 | 0.0117 | 0.0219 | 0.0540 | 0.1196 |
| | | TIME 6HR | 7.85 | 0.0243 | 0.0873 | -0.0004 | 0.0483 | 0.0025 | 0.0169 | 0.0020 | 0.0300 | 0.0403 | 0.0742 |
| | | TIME 12HR | 6.31 | 0.1414 | 0.0652 | 0.0094 | 0.0246 | -0.0139 | 0.0170 | -0.0217 | 0.0215 | -0.0789 | 0.0583 |
| | | TIME 12HR | 6.88 | 0.1201 | 0.0587 | -0.0742 | 0.0363 | -0.0020 | 0.0310 | -0.0599 | 0.0277 | 0.0345 | 0.0595 |
| | | TIME 24HR | 5.89 | 0.2293 | 0.0537 | -0.0421 | 0.0304 | 0.0006 | 0.0307 | -0.0183 | 0.0275 | -0.0700 | 0.0607 |
| | | TIME 24HR | 5.81 | 0.2287 | 0.0251 | -0.0841 | 0.0359 | -0.0004 | 0.0313 | -0.0124 | 0.0276 | -0.0093 | 0.0602 |
| | | TIME 48HR | 3.87 | 0.3874 | 0.0183 | -0.0128 | 0.0302 | -0.0146 | 0.0308 | -0.0298 | 0.0275 | -0.1125 | 0.0614 |
| | | TIME 48HR | 5.18 | 0.3584 | 0.0204 | -0.0449 | 0.0236 | -0.0279 | 0.0189 | -0.0695 | 0.0182 | -0.0292 | 0.0518 |
| Chemo. | Sterile control #1 | MED. ctl | 8.70 | 0.2617 | 0.0225 | -0.0184 | 0.0155 | 0.0229 | 0.0117 | 0.0286 | 0.0303 | 0.1104 | 0.0296 |
| | | H2O ctl | 10.03 | 0.0511 | 0.0244 | 0.0647 | 0.0136 | 0.0652 | 0.0164 | 0.0081 | 0.0338 | 0.0217 | 0.0242 |
| | | TIME 0HR | 9.29 | -0.0087 | 0.0322 | 0.0168 | 0.0366 | 0.0312 | 0.0132 | 0.0052 | 0.0298 | 0.0028 | 0.0521 |
| | | TIME 0HR | 7.97 | 0.1609 | 0.0328 | 0.0029 | 0.0335 | 0.0576 | 0.0146 | -0.0314 | 0.0314 | 0.0847 | 0.0403 |
| | | TIME 3HR | 8.39 | 0.0734 | 0.0207 | -0.0161 | 0.0375 | -0.0012 | 0.0150 | 0.0333 | 0.0246 | 0.0304 | 0.0547 |
| | | TIME 3HR | 8.67 | 0.0585 | 0.0221 | -0.0083 | 0.0373 | 0.0204 | 0.0153 | 0.0147 | 0.0266 | -0.0294 | 0.0627 |
| | | TIME 6HR | 8.05 | 0.0618 | 0.0117 | 0.0412 | 0.0110 | 0.0219 | 0.0148 | -0.0060 | 0.0240 | -0.0075 | 0.0733 |
| | | TIME 6HR | 7.88 | 0.1531 | 0.0243 | 0.0056 | 0.0364 | -0.0019 | 0.0157 | -0.0465 | 0.0165 | -0.0101 | 0.0744 |
| | | TIME 12HR | 7.66 | 0.1457 | 0.0258 | 0.0404 | 0.0239 | -0.0247 | 0.0339 | -0.0044 | 0.0169 | -0.0710 | 0.0419 |
| | | TIME 12HR | 7.73 | 0.1354 | 0.0245 | 0.0466 | 0.0272 | 0.0521 | 0.0251 | 0.0280 | 0.0180 | -0.0188 | 0.0453 |
| | | TIME 24HR | 8.01 | 0.1056 | 0.0372 | -0.0422 | 0.0271 | -0.0213 | 0.0072 | -0.0110 | 0.0252 | 0.0176 | 0.0511 |
| | | TIME 24HR | 7.89 | 0.1688 | 0.0381 | -0.0081 | 0.0224 | 0.0218 | 0.0113 | 0.0135 | 0.0270 | -0.0151 | 0.0527 |
| | | TIME 48HR | 7.84 | 0.1197 | 0.0223 | -0.0301 | 0.0187 | -0.0096 | 0.0265 | -0.0108 | 0.0135 | 0.0449 | 0.0148 |
| | | TIME 48HR | 7.64 | 0.1428 | 0.0226 | -0.0343 | 0.0181 | -0.0572 | 0.0243 | -0.0111 | 0.0142 | 0.0197 | 0.0291 |

Continued on next page

**Table S1 (continued)**: Total Hg concentrations and Hg isotope ratios for the reactant pool $Hg^{II}$ in phototrophic and chemotrophic growth conditions experiments. Each line corresponds to an individual sample and technical duplicates were taken for each time point. Std represents the standard deviation from analytical replicates on the MC-ICP-MS. Abbreviations denote Metabo. = Metabolism; Photo. = Phototrophy; Chemo. = Chemotrophy; Treat. = Treatment; Hg conc. = Hg concentration, rep = replicate; Avg = average; Std = Standard deviation; ctl = control, MED = medium. $\delta^{202}$Hg values in **bold** denote those that deviate substantially from the composition of the NIST 3133 standard in medium spiked controls.

| Metabo. | Treat. | Sample | Hg conc. (nM) | $\delta^{202}$Hg Avg | $\delta^{202}$Hg Std | $\Delta^{199}$Hg Avg | $\Delta^{199}$Hg Std | $\Delta^{200}$Hg Avg | $\Delta^{200}$Hg Std | $\Delta^{201}$Hg Avg | $\Delta^{201}$Hg Std | $\Delta^{204}$Hg Avg | $\Delta^{204}$Hg Std |
|---|---|---|---|---|---|---|---|---|---|---|---|---|---|
| Chemo. | Sterile ctl #2 | MED. Blk | 5.96 | **0.5452** | 0.0224 | -0.0685 | 0.0232 | -0.0212 | 0.0238 | -0.0402 | 0.0248 | -0.0023 | 0.0256 |
| | | H2O Blk | 9.54 | -0.0420 | 0.0114 | -0.0508 | 0.0234 | -0.0284 | 0.0359 | -0.0154 | 0.0288 | 0.0647 | 0.0183 |
| | | TIME 0HR | 7.31 | 0.1773 | 0.0469 | 0.0073 | 0.0329 | 0.0324 | 0.0324 | -0.0209 | 0.0163 | -0.0355 | 0.0431 |
| | | TIME 0HR | 7.25 | 0.0331 | 0.0073 | -0.0348 | 0.0322 | -0.0045 | 0.0600 | -0.0308 | 0.0484 | 0.0048 | 0.0096 |
| | | TIME 3HR | 7.26 | 0.1694 | 0.0233 | -0.0121 | 0.0234 | 0.0164 | 0.0241 | -0.0399 | 0.0141 | -0.0422 | 0.0665 |
| | | TIME 3HR | 7.70 | 0.2087 | 0.0268 | -0.0167 | 0.0215 | 0.0289 | 0.0300 | -0.0263 | 0.0138 | 0.0368 | 0.0663 |
| | | TIME 6HR | 7.70 | 0.1382 | 0.0677 | 0.0066 | 0.0196 | 0.0072 | 0.0060 | 0.0067 | 0.0154 | -0.0164 | 0.0688 |
| | | TIME 6HR | 7.63 | 0.0607 | 0.0398 | -0.0030 | 0.0399 | -0.0204 | 0.0417 | -0.0265 | 0.0085 | 0.0057 | 0.0715 |
| | | TIME 12HR | 7.64 | 0.2430 | 0.0340 | 0.0205 | 0.0469 | 0.0273 | 0.0466 | 0.0022 | 0.0129 | -0.0111 | 0.0205 |
| | | TIME 12HR | 7.37 | 0.1662 | 0.0389 | -0.0103 | 0.0470 | 0.0055 | 0.0480 | -0.0301 | 0.0053 | -0.0656 | 0.0216 |
| | | TIME 24HR | 5.15 | 0.8339 | 0.0211 | -0.1161 | 0.0395 | -0.0059 | 0.0435 | -0.0548 | 0.0119 | -0.0923 | 0.0229 |
| | | TIME 24HR | 7.52 | 0.1225 | 0.0205 | -0.0668 | 0.0448 | 0.0030 | 0.0403 | -0.0165 | 0.0326 | -0.0203 | 0.0174 |
| | | TIME 48HR | 7.25 | 0.1940 | 0.0222 | -0.0176 | 0.0435 | 0.0070 | 0.0417 | -0.0300 | 0.0174 | -0.0097 | 0.0354 |
| | | TIME 48HR | 7.63 | 0.1408 | 0.0108 | -0.0243 | 0.0276 | 0.0040 | 0.0251 | 0.0008 | 0.0317 | -0.0227 | 0.0353 |
| Chemo. | Live cells rep #1 | MED. ctl | 3.18 | NA | NA | NA | NA | NA | NA | NA | NA | NA | NA |
| | | H2O Blkctl | 10.90 | 0.0384 | 0.0068 | -0.0507 | 0.0278 | 0.0298 | 0.0125 | 0.0229 | 0.0234 | 0.0383 | 0.0282 |
| | | TIME 0HR | 8.01 | 0.0726 | 0.0099 | -0.0730 | 0.0204 | -0.0215 | 0.0175 | -0.0334 | 0.0213 | -0.0296 | 0.0481 |
| | | TIME 0HR | 8.09 | 0.1379 | 0.0171 | -0.0187 | 0.0363 | -0.0395 | 0.0079 | -0.0096 | 0.0228 | -0.0136 | 0.0417 |
| | | TIME 3HR | 6.08 | -0.2680 | 0.0200 | -0.0418 | 0.0374 | -0.0347 | 0.0136 | 0.0045 | 0.0200 | 0.0351 | 0.0416 |
| | | TIME 3HR | 5.53 | -0.2759 | 0.0100 | -0.0286 | 0.0382 | -0.0008 | 0.0092 | -0.0003 | 0.0211 | -0.0034 | 0.0409 |
| | | TIME 6HR | 5.68 | -0.2896 | 0.0107 | -0.0055 | 0.0377 | -0.0144 | 0.0061 | -0.0272 | 0.0075 | -0.0533 | 0.0515 |
| | | TIME 6HR | 5.39 | -0.2237 | 0.0113 | -0.0050 | 0.0136 | -0.0011 | 0.0323 | -0.0396 | 0.0233 | -0.0545 | 0.0369 |
| | | TIME 12HR | 5.27 | -0.1774 | 0.0162 | -0.0642 | 0.0254 | -0.0120 | 0.0358 | -0.0264 | 0.0344 | -0.0024 | 0.0198 |
| | | TIME 12HR | 5.19 | -0.1756 | 0.0099 | -0.0439 | 0.0132 | -0.0134 | 0.0391 | 0.0026 | 0.0246 | -0.0787 | 0.0493 |
| | | TIME 24HR | 4.51 | 0.1089 | 0.0089 | -0.0386 | 0.0200 | 0.0019 | 0.0398 | 0.0126 | 0.0269 | -0.0876 | 0.0400 |
| | | TIME 24HR | 4.59 | 0.0276 | 0.0232 | -0.0509 | 0.0128 | -0.0086 | 0.0428 | -0.0356 | 0.0247 | -0.1025 | 0.0294 |
| | | TIME 48HR | 3.59 | 0.1492 | 0.0380 | -0.0607 | 0.0220 | 0.0413 | 0.0207 | 0.0215 | 0.0292 | 0.1010 | 0.0329 |
| | | TIME 48HR | 3.66 | 0.1913 | 0.0389 | -0.0256 | 0.0275 | 0.0179 | 0.0145 | -0.0323 | 0.0294 | -0.0428 | 0.0340 |

Continued on next page

**Table S1 (continued)**: Total Hg concentrations and Hg isotope ratios for the reactant pool $Hg^{II}$ in phototrophic and chemotrophic growth conditions experiments. Each line corresponds to an individual sample and technical duplicates were taken for each time point. Std represents the standard deviation from analytical replicates on the MC-ICP-MS. Abbreviations denote Metabo. = Metabolism; Photo. = Phototrophy; Chemo. = Chemotrophy; Treat. = Treatment; Hg conc. = Hg concentration, rep = replicate; Avg = average; Std = Standard deviation; ctl = control, MED = medium. $\delta^{202}$Hg values in **bold** denote those that deviate substantially from the composition of the NIST 3133 standard in medium spiked controls.

| Metabo. | Treat. | Sample | Hg conc. (nM) | $\delta^{202}$Hg Avg | $\delta^{202}$Hg Std | $\Delta^{199}$Hg Avg | $\Delta^{199}$Hg Std | $\Delta^{200}$Hg Avg | $\Delta^{200}$Hg Std | $\Delta^{201}$Hg Avg | $\Delta^{201}$Hg Std | $\Delta^{204}$Hg Avg | $\Delta^{204}$Hg Std |
|---|---|---|---|---|---|---|---|---|---|---|---|---|---|
| Chemo. | Live cells rep #2 | MED. ctl | 5.80 | **0.5401** | 0.0196 | -0.0043 | 0.0277 | 0.0484 | 0.0251 | 0.0183 | 0.0142 | 0.0241 | 0.0384 |
| | | H2O ctl | 9.92 | 0.0594 | 0.0186 | 0.0400 | 0.0280 | -0.0064 | 0.0080 | -0.0122 | 0.0274 | -0.0232 | 0.0096 |
| | | TIME 0HR | 8.10 | 0.1632 | 0.0225 | -0.0264 | 0.0300 | 0.0188 | 0.0328 | -0.0341 | 0.0173 | -0.0369 | 0.0430 |
| | | TIME 0HR | 7.98 | 0.0387 | 0.0249 | -0.0234 | 0.0320 | 0.0108 | 0.0311 | 0.0052 | 0.0195 | 0.0311 | 0.0404 |
| | | TIME 3HR | 7.44 | 0.2343 | 0.0220 | 0.0031 | 0.0327 | 0.0221 | 0.0078 | -0.0318 | 0.0222 | -0.0303 | 0.0328 |
| | | TIME 3HR | 7.63 | 0.2620 | 0.0225 | -0.0092 | 0.0300 | -0.0140 | 0.0328 | -0.0503 | 0.0173 | -0.0625 | 0.0430 |
| | | TIME 6HR | 6.99 | 0.1904 | 0.0083 | 0.0071 | 0.0347 | 0.0111 | 0.0291 | -0.0305 | 0.0232 | -0.0721 | 0.0412 |
| | | TIME 6HR | 7.09 | 0.1580 | 0.0200 | -0.0055 | 0.0310 | 0.0307 | 0.0317 | -0.0110 | 0.0133 | -0.0246 | 0.0532 |
| | | TIME 12HR | 6.92 | 0.1740 | 0.0343 | -0.0336 | 0.0148 | -0.0152 | 0.0255 | -0.0410 | 0.0196 | -0.0296 | 0.0503 |
| | | TIME 12HR | 6.58 | 0.1377 | 0.0205 | -0.0424 | 0.0258 | -0.0229 | 0.0234 | -0.0081 | 0.0202 | -0.1049 | 0.0475 |
| | | TIME 24HR | 3.69 | 0.7127 | 0.0376 | -0.0753 | 0.0170 | -0.0059 | 0.0115 | -0.0328 | 0.0314 | -0.0167 | 0.0259 |
| | | TIME 24HR | 3.59 | 0.7185 | 0.0399 | -0.0515 | 0.0256 | -0.0136 | 0.0115 | -0.0578 | 0.0348 | -0.0464 | 0.0177 |
| | | TIME 48HR | 0.47 | NA | NA | NA | NA | NA | NA | NA | NA | NA | NA |
| | | TIME 48HR | 3.14 | NA | NA | NA | NA | NA | NA | NA | NA | NA | NA |

**Table S2:** Total Hg concentrations and Hg isotope ratios for the $Hg^0$ pool in phototrophic and chemotrophic growth conditions experiments. Each line corresponds to the Hg present on two gold traps used to sample each time point in parallel. Std represents the standard deviation from analytical replicates on the MC-ICP-MS. Abbreviations denote the following: Metabo. = Metabolism; Photo. = Phototrophy; Chemo. = Chemotrophy; Treat. = Treatment; Hg conc. = Hg concentration, rep = replicate; Avg = average; Std = Standard deviation.

| Metabo. | Treat. | Sample | Hg on trap (ng) | $\delta^{202}$Hg Avg | $\delta^{202}$Hg Std | $\Delta^{199}$Hg Avg | $\Delta^{199}$Hg Std | $\Delta^{200}$Hg Avg | $\Delta^{200}$Hg Std | $\Delta^{201}$Hg Avg | $\Delta^{201}$Hg Std | $\Delta^{204}$Hg Avg | $\Delta^{204}$Hg Std |
|---|---|---|---|---|---|---|---|---|---|---|---|---|---|
| Photo. | Sterile control | TIME 3HR | 0.0137 | NA | NA | NA | NA | NA | NA | NA | NA | NA | NA |
| | | TIME 6HR | 0.0036 | NA | NA | NA | NA | NA | NA | NA | NA | NA | NA |
| | | TIME 12HR | 0.0221 | NA | NA | NA | NA | NA | NA | NA | NA | NA | NA |
| | | TIME 24HR | 0.0030 | NA | NA | NA | NA | NA | NA | NA | NA | NA | NA |
| | | TIME 48HR | 0.0037 | NA | NA | NA | NA | NA | NA | NA | NA | NA | NA |
| | Live cells rep #1 | TIME 3HR | 0.1256 | 0.50 | 0.06 | 0.16 | 0.05 | -0.01 | 0.03 | 0.15 | 0.03 | -0.12 | 0.10 |
| | | TIME 6HR | 0.1484 | -0.98 | 0.05 | 0.05 | 0.06 | -0.01 | 0.03 | 0.02 | 0.03 | -0.01 | 0.02 |
| | | TIME 12HR | 0.2254 | -0.74 | 0.05 | 0.12 | 0.06 | 0.02 | 0.03 | 0.12 | 0.04 | -0.03 | 0.06 |
| | | TIME 24HR | 0.2187 | -0.49 | 0.03 | 0.15 | 0.05 | 0.04 | 0.02 | 0.10 | 0.02 | -0.11 | 0.07 |
| | | TIME 48HR | 0.0896 | -0.04 | 0.02 | 0.15 | 0.01 | -0.01 | 0.01 | 0.03 | 0.03 | -0.16 | 0.06 |
| | Live cells rep #2 | TIME 3HR | 0.1064 | 0.66 | 0.04 | 0.17 | 0.05 | -0.03 | 0.02 | 0.06 | 0.03 | -0.02 | 0.07 |
| | | TIME 6HR | 0.1155 | -0.81 | 0.06 | 0.10 | 0.03 | -0.02 | 0.01 | 0.04 | 0.03 | 0.06 | 0.11 |
| | | TIME 12HR | 0.1487 | -0.69 | 0.07 | 0.21 | 0.05 | 0.00 | 0.03 | 0.11 | 0.02 | -0.04 | 0.13 |
| | | TIME 24HR | 0.1520 | -0.34 | 0.07 | 0.17 | 0.02 | 0.08 | 0.02 | 0.11 | 0.01 | -0.04 | 0.05 |
| | | TIME 48HR | 0.0805 | 0.04 | 0.07 | 0.15 | 0.02 | 0.02 | 0.02 | 0.03 | 0.04 | -0.13 | 0.08 |

Continued on next page

**Table S2 (continued):** Total Hg concentrations and Hg isotope ratios for the $Hg^0$ pool in phototrophic and chemotrophic growth conditions experiments. Each line corresponds to the Hg present on two gold traps used to sample each time point in parallel. Std represents the standard deviation from analytical replicates on the MC-ICP-MS. Abbreviations denote the following: Metabo. = Metabolism; Photo. = Phototrophy; Chemo. = Chemotrophy; Treat. = Treatment; Hg conc. = Hg concentration, rep = replicate; Avg = average; Std = Standard deviation.

| Metabo. | Treat. | Sample | Hg on trap (ng) | δ202Hg Avg | δ202Hg Std | Δ199Hg Avg | Δ199Hg Std | Δ200Hg Avg | Δ200Hg Std | Δ201Hg Avg | Δ201Hg Std | Δ204Hg Avg | Δ204Hg Std |
|---|---|---|---|---|---|---|---|---|---|---|---|---|---|
| Chemo. | Sterile control #1 | TIME 3HR | 0.0256 | NA | NA | NA | NA | NA | NA | NA | NA | NA | NA |
| | | TIME 6HR | 0.0020 | NA | NA | NA | NA | NA | NA | NA | NA | NA | NA |
| | | TIME 12HR | 0.0045 | NA | NA | NA | NA | NA | NA | NA | NA | NA | NA |
| | | TIME 24HR | 0.0040 | NA | NA | NA | NA | NA | NA | NA | NA | NA | NA |
| | | TIME 48HR | 0.0015 | NA | NA | NA | NA | NA | NA | NA | NA | NA | NA |
| . | Sterile control #2 | TIME 3HR | 0.0085 | NA | NA | NA | NA | NA | NA | NA | NA | NA | NA |
| | | TIME 6HR | 0.0013 | NA | NA | NA | NA | NA | NA | NA | NA | NA | NA |
| | | TIME 12HR | 0.0040 | NA | NA | NA | NA | NA | NA | NA | NA | NA | NA |
| | | TIME 24HR | 0.0021 | NA | NA | NA | NA | NA | NA | NA | NA | NA | NA |
| | | TIME 48HR | 0.0010 | NA | NA | NA | NA | NA | NA | NA | NA | NA | NA |
| | Live cells rep #1 | TIME 3HR | 0.6376 | 0.89 | 0.02 | 0.20 | 0.03 | 0.02 | 0.02 | 0.17 | 0.02 | -0.02 | 0.03 |
| | | TIME 6HR | 0.1542 | 0.90 | 0.03 | 0.08 | 0.03 | -0.01 | 0.01 | 0.06 | 0.02 | 0.03 | 0.03 |
| | | TIME 12HR | 0.3673 | NA | NA | NA | NA | NA | NA | NA | NA | NA | NA |
| | | TIME 24HR | 0.1662 | -1.79 | 0.02 | 0.19 | 0.02 | 0.03 | 0.02 | 0.15 | 0.01 | -0.05 | 0.05 |
| | | TIME 48HR | 0.3141 | -0.59 | 0.03 | 0.17 | 0.02 | 0.04 | 0.01 | 0.12 | 0.02 | -0.02 | 0.05 |
| | Live cells rep #2 | TIME 3HR | 0.0002 | NA | NA | NA | NA | NA | NA | NA | NA | NA | NA |
| | | TIME 6HR | 0.0133 | NA | NA | NA | NA | NA | NA | NA | NA | NA | NA |
| | | TIME 12HR | 0.1642 | -0.91 | 0.01 | 0.17 | 0.02 | 0.03 | 0.01 | 0.11 | 0.02 | 0.02 | 0.05 |
| | | TIME 24HR | 1.0858 | 0.27 | 0.01 | 0.07 | 0.02 | -0.02 | 0.01 | 0.07 | 0.01 | -0.03 | 0.03 |
| | | TIME 48HR | 0.0627 | 1.73 | 0.01 | 0.10 | 0.01 | 0.02 | 0.01 | 0.09 | 0.01 | -0.03 | 0.04 |

**Table S3:** Summary of regression models for mass dependent fractionation of $^{202}Hg^{II}$ and $^{202}Hg^{0}$ in phototrophically and chemotrophically grown *H. modesticaldum* cultures. P-values for regressions that displayed a significant ($p<0.05$) fit are shown in **bold**.

| | Phototrophic | | | | Chemotrophic | | | |
|---|---|---|---|---|---|---|---|---|
| | $Hg^{II}$ model | | $Hg^{0}$ model | | $Hg^{II}$ model | | $Hg^{0}$ model | |
| | Replicate 1 | Replicate 2 | Replicate 1 | Replicate 2 | Replicate 1 | Replicate 2 | Replicate 1 | Replicate 2 |
| Slope | -0.00100 | -0.00060 | -0.00380 | -0.00192 | -0.00132 | -0.00095 | 0.00538 | -0.00434 |
| Intercept | $-5.66 \times 10^{-5}$ | $-8.85 \times 10^{-5}$ | -0.002 | -0.0018 | -0.0008 | $-1.19 \times 10^{-5}$ | 0.0016 | $-7.23 \times 10^{-5}$ |
| Adjusted $R^2$ | 0.8496 | 0.8246 | 0.9286 | 0.8979 | 0.9275 | 0.8925 | 0.2278 | 0.7008 |
| p-value | **0.0057** | **0.0078** | **0.0241** | **0.0346** | **0.0055** | **0.01** | 0.3034 | 0.2528 |
| $\varepsilon_{202/198}$ (‰) | -1.00 | -0.60 | -3.80 | -1.92 | -1.32 | -0.95 | NA | NA |

**Table S4:** Comparison of measured and calculated isotope fractionation values for the reactant pool of $Hg^{II}$ using a mass balance approach. Measured values of Hg concentration and $\delta^{202}Hg$ are represented as an average of replicates from experiments presented in Tables S1 and S3. Predicted values for media were obtained using the equations outlined for mass balance calculations. Differences between the measured and predicted values that are substantially greater than the analytical variability of measurements (2SD = 0.08) are shown in **bold**. Abbreviations denote Photo. = phototrophic, Chemo. = chemotrophic, rep = replicates, Avg. = average, Pred. = predicted.

| Metabo. | Treat. | Sample | Avg THg (nM) | Avg $\delta^{202}$Hg-Media | Avg f $Hg^{II}$ | Avg. $\delta^{202}$Hg-$Hg^0$ | Pred. $\delta^{202}$Hg-media | Difference (measured - predicted) |
|---|---|---|---|---|---|---|---|---|
| Photo. | Live cells rep #1 | TIME 0HR | 9.49 | 0.03105 | 1 | NA | NA | NA |
| | | TIME 3HR | 8.95 | 0.0452 | 0.9433674 | 0.5 | -0.0300 | 0.0752 |
| | | TIME 6HR | 8.26 | 0.04405 | 0.86977136 | -0.98 | 0.1467 | -0.1027 |
| | | TIME 12HR | 7.82 | 0.0979 | 0.82362238 | -0.74 | 0.1585 | -0.0606 |
| | | TIME 24HR | 6.82 | 0.29345 | 0.71820672 | -0.49 | 0.1923 | 0.1012 |
| | | TIME 48HR | 6.26 | 0.37145 | 0.65915077 | -0.04 | 0.0207 | **0.3508** |
| Photo. | Live cells rep #2 | TIME 0HR | 9.07 | 0.0856 | 1 | NA | NA | NA |
| | | TIME 3HR | 8.53 | -0.0214 | 0.94040465 | 0.66 | -0.0418 | 0.0204 |
| | | TIME 6HR | 8.10 | 0.0412 | 0.89282761 | -0.81 | 0.0972 | -0.0560 |
| | | TIME 12HR | 6.60 | 0.13075 | 0.72743812 | -0.69 | 0.2585 | -0.1278 |
| | | TIME 24HR | 5.85 | 0.229 | 0.64507415 | -0.34 | 0.1871 | 0.0419 |
| | | TIME 48HR | 4.52 | 0.3729 | 0.49864932 | 0.04 | -0.0402 | **0.4131** |
| Chemo. | Live cells rep #1 | TIME 0HR | 8.05 | 0.10525 | 1 | NA | NA | NA |
| | | TIME 3HR | 5.80 | -0.27195 | 0.72101112 | 0.89 | -0.3444 | 0.0724 |
| | | TIME 6HR | 5.54 | -0.25665 | 0.68784548 | 0.9 | -0.4084 | 0.1518 |
| | | TIME 12HR | 5.23 | -0.1765 | 0.64933855 | NA | NA | NA |
| | | TIME 24HR | 4.55 | 0.06825 | 0.56524439 | -1.79 | 1.3768 | **-1.3085** |
| | | TIME 48HR | 3.63 | 0.17025 | 0.45046891 | -0.59 | 0.7197 | **-0.5495** |
| Chemo. | Live cells rep #2 | TIME 0HR | 8.04 | 0.10095 | 1 | NA | NA | NA |
| | | TIME 3HR | 7.53 | 0.24815 | 0.93726295 | NA | NA | NA |
| | | TIME 6HR | 7.04 | 0.1742 | 0.87558291 | NA | NA | NA |
| | | TIME 12HR | 6.75 | 0.15585 | 0.83939564 | -0.91 | 0.1741 | -0.0183 |
| | | TIME 24HR | 3.64 | 0.7156 | 0.45271405 | 0.27 | -0.3264 | **1.0420** |
| | | TIME 48HR | 1.81 | NA | NA | 1.73 | NA | NA |

**Table S5:** Comparison of enrichment factors and measures of variability used in this study and the original cited work. Values in bold represent calculation ratios that differ between the original article cited and the current study. Abbreviations: SD = standard deviation, SE = standard error, p = product, r = reactant, Ox. = oxygenic, Anox. = anoxygenic, PS = photosynthesis.

| Hg transformation | Pathway | Microorganism or treatment | $\varepsilon_{p/r}$ value used (‰) | 2SD from this work | Original ε value (‰) | Original ε ratio | Original α value | Original α ratio | Variability value | SD vs SE | Comments | Ref. |
|---|---|---|---|---|---|---|---|---|---|---|---|---|
| Reduction | Abiotic photoreduction | Low DOC natural sunlight | -0.6 | 0.282 | NA | NA | 0.9994 | **p/r** | 0.0002 | 2SE | Experiments were conducted in replicates of n =2, which was used to convert SE to 2SD. Converted 2SD for α to 2SD for ε by multiplying by 1000. | (3) |
| Reduction | Abiotic photoreduction | Marine exudate low UVB | -1.47 | NA | 1.47 | r/p | NA | r/p | NA | NA | Calculated new ε value by taking reciprocal. No alpha values provided in supplementary table where ε was obtained. | (4) |
| Reduction | Ox. PS. reduction | *Isochrysis galbana* high UV | -0.14 | NA | 0.14 | r/p | NA | r/p | NA | NA | Same as above. | (4) |
| Reduction | Ox. PS. reduction | *Isochrysis galbana* low UV | -0.61 | NA | 0.61 | r/p | NA | r/p | NA | NA | Same as above. | (4) |
| Reduction | Aerobic MerA reduction | *Escherichia coli* JM109/pPB117 | -1.4 | 0.1 | NA | NA | 1.0014 | r/p | 0.0001 | 2SD | Converted 2SD for α to 2SD for ε by multiplying by 1000. | (2) |
| Reduction | Aerobic MerA reduction | *Bacillus cereus* strain 5 | -1.2 | 0.1 | NA | NA | 1.0012 | r/p | 0.0001 | 2SD | Converted 2SD for α to 2SD for ε by multiplying by 1000. | (1) |
| Reduction | Aerobic MerA reduction | *Anoxybacillus* sp. FB9 | -1.4 | 0.1 | NA | NA | 1.0014 | r/p | 0.0001 | 2SD | Same as above. | (1) |
| Reduction | Anox. PS. reduction | *Heliobacterium modesticaldum* Ice1 | -0.801 | 0.561 | NA | NA | NA | NA | NA | NA | See methods | This study |
| Reduction | Anaerobic respiratory reduction | *Shewanella oneidensis* MR-1 | -1.8 | 0.3 | NA | NA | 1.0018 | r/p | 0.0003 | 2SD | Converted 2SD for α to 2SD for ε by multiplying by 1000. | (1) |
| Reduction | Anaerobic respiratory reduction | *Geobacter sulfurreducens* PCA ΔhgcAB | -1.6 | NA | NA | NA | 1.0016 | r/p | | | Values from regression analysis in original article supplementary material | (5) |
| Oxidation | Abiotic dark oxidation | L-cysteine | 1.51 | 0.2 | 1.51 | p/r | NA | NA | 0.05 | 2SE | Uncertainty was derived from analytical measures of reference material. Based on the sample size presented in supplementary information for reference material, n =4 was used to convert SE to 2SD. | (6) |
| Oxidation | Abiotic dark oxidation | Glutathione | 1.47 | 0.24 | 1.47 | p/r | NA | NA | 0.06 | 2SE | Same as above. | (6) |
| Methylation | Anaerobic respiratory methylation | *Geobacter sulfurreducens* PCA | -0.92 | NA | -0.92 | p/r | 1.0009 | r/p | NA | NA | Values verified directly in regression analyses provided in original article's supplementary material. | (7) |
| Methylation | Anaerobic respiratory methylation | *Desulfovibrio desulfuricans* ND132 | -1.1 | NA | -1.1 | p/r | 1.0011 | r/p | NA | NA | Same as above. | (7) |
| Methylation | Anaerobic respiratory methylation (sulphate reduction) | *Desulfovibrio dechloracetivorans* BerOc1 | -2.5 | 1.9 | NA | NA | 1.0025 | r/p | 0.0011 | Max SE | Authors applied max SE from all experiments to measure variability. This value is compared to other studies as SE but the values actually represent 2SD. Supplementary information suggests n = 3 experiments were performed, which was used to convert SE to 2SD. Converted 2SD for α to 2SD for ε by multiplying by 1000. | (8)[a] |
| Methylation | Anaerobic respiratory methylation (fumarate reduction | *Desulfovibrio dechloracetivorans* BerOc1 | -4.4 | 1.9 | NA | NA | 1.0044 | r/p | 0.0011 | Max SE | Same as above. | (8)[a] |
| Methylation | Fermentative methylation | *Desulfobulbus propionicus* MUD10 | -2.6 | -0.4 | NA | NA | 1.0026 | r/p | 0.0004 | 2SD | Converted 2SD for α to 2SD for ε by multiplying by 1000. | (9)[a] |
| Reduction | Fermentative reduction | *Heliobacterium modesticaldum* Ice1 | -1.137 | 0.515 | NA | NA | NA | NA | NA | NA | See methods | This study |

a. These values were not included in the main text because they were obtained with a MC-ICPM-MS method that uses gas chromatography and differs substantially from the methods used in other studies compiled here.
